# Exenatide administration time determines the effects on blood pressure dipping in *db/db* mice via modulation of food intake and sympathetic activity

**DOI:** 10.1101/2024.07.02.601700

**Authors:** Aaron N. Chacon, Wen Su, Tianfei Hou, Zhenheng Guo, Ming C. Gong

## Abstract

Type 2 diabetics have an increased prevalence of hypertension and nondipping blood pressure (BP), which worsen cardiovascular outcomes. Exenatide, a short acting glucagon-like peptide-1 receptor agonist (GLP-1RA) used to treat type 2 diabetes, also demonstrates blood pressure (BP)-lowering effects. However, the mechanisms behind this and the impact of administration timing on BP dipping remain unclear. We investigated the effects of exenatide intraperitoneal injected at light onset (ZT0) or dark onset (ZT12) in diabetic (db/db) mice and nondiabetic controls. Using radio-telemetry and BioDAQ cages, we continuously monitored BP and food intake. Db/db mice exhibited non-dipping BP and increased food intake. ZT0 exenatide administration restored BP dipping by specifically lowering light-phase BP, while ZT12 exenatide reversed dipping by lowering dark-phase BP. These effects correlated with altered food intake patterns, and importantly, were abolished when food access was removed. Additionally, urinary norepinephrine excretion, measured by HPLC, was significantly reduced 6 hours post-exenatide at both ZT0 and ZT12, suggesting sympathetic nervous system involvement. Notably, combining exenatide with either ganglionic blocker mecamylamine or α-blocker prazosin did not enhance BP reduction beyond the individual effects of each blocker. These findings reveal that exenatide, when administered at light onset, restores BP dipping in db/db mice by suppressing light-phase food intake and sympathetic activity. Importantly, the efficacy of exenatide is dependent on food availability and its timing relative to circadian rhythms, highlighting the potential for chronotherapy in optimizing GLP-1RA- based treatments for type 2 diabetes and hypertension.

**Graphic Abstract:** 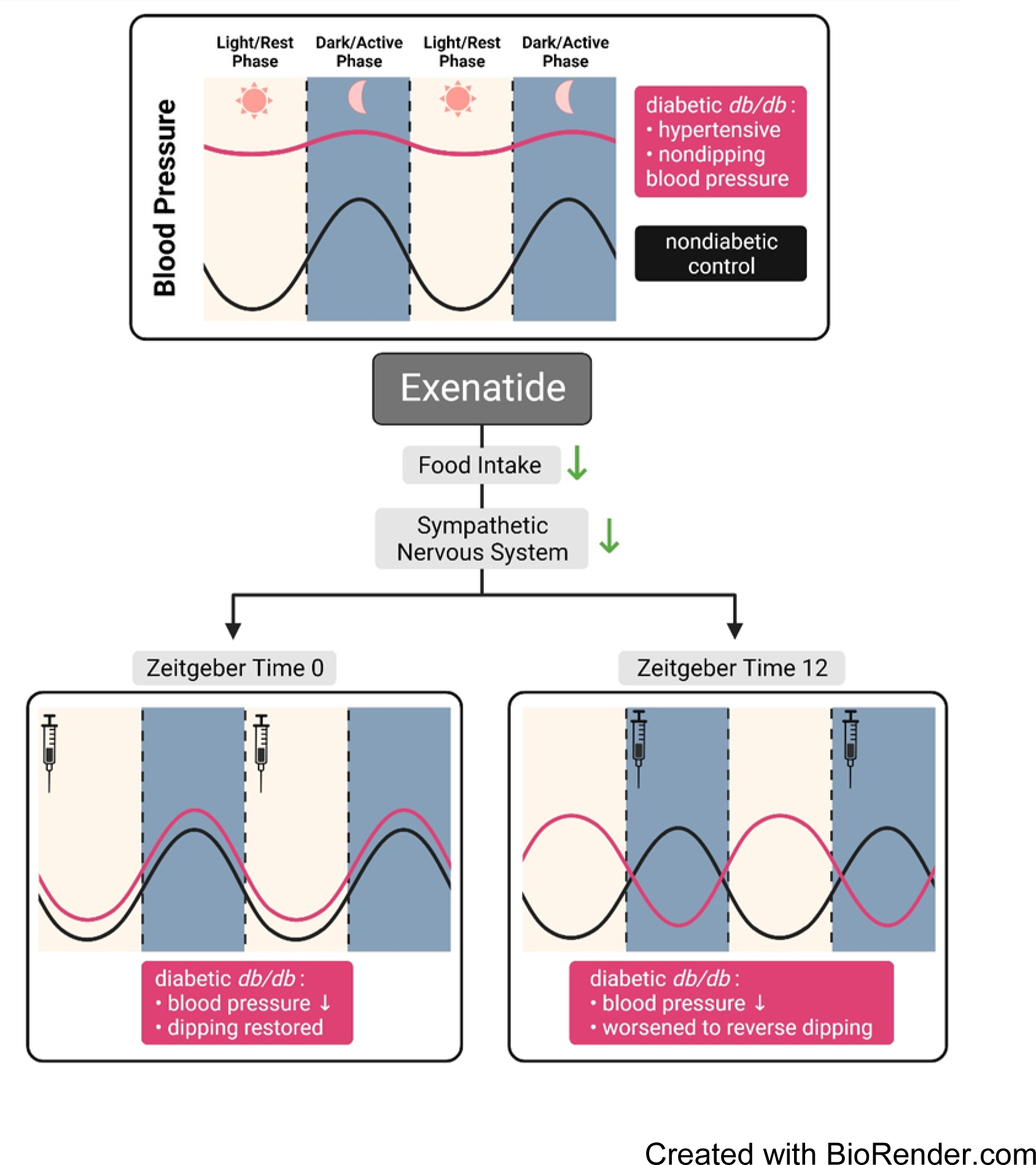

**Article Highlights:**

- Maintaining a normal blood pressure (BP) circadian rhythm is vital for cardiovascular health, but diabetes often disrupts this rhythm. The effect of exenatide, a GLP-1 receptor agonist (GLP-1RA), on BP rhythm in diabetes is uncertain.
- This study investigates the impact of exenatide administration timing on BP patterns in diabetic db/db mice.
- Findings indicate that exenatide given at the onset of rest restores normal BP dipping, while at the start of the active phase worsens BP rhythm by modulating food intake and sympathetic activity.
- Timing GLP-1 RA administration may optimize BP control and provide cardiovascular benefits for type 2 diabetes patients.

## Introduction

Type 2 diabetes (T2DM) is an escalating global concern, affecting approximately 383 million individuals worldwide, and is projected to surge to nearly 600 million cases by 2035^7^. Cardiovascular complications are principally responsible for increased morbidity and mortality in type 2 diabetics. Of particular relevance, hypertension has shown an elevated prevalence in those with T2DM and has been identified as a prominent risk factor for detrimental cardiovascular events within this cohort^8^. T2DM and cardiovascular disease are inextricably linked, with T2DM patients exhibiting a substantially higher risk of coronary artery disease, myocardial infarction^9^, and cardiovascular mortality^10,11^, attributable to a high prevalence of hypertension^12,13^ in this population. Additionally, emerging evidence underscores the increased incidence of nondipping blood pressure (BP) in diabetic subjects – defined by a reduction in BP of less than 10% during nighttime hours – compared to non-diabetic counterparts^14,15^. Nondipping BP has been associated with increased cardiovascular risk^16,17^ and mortality^18^, lower rates of creatine clearance^19^, a decline in glomerular filtration rate^20^, and potentially target organ damage cardiac damage^16,21,22^. As such, the effective management of hypertension and nondipping BP is paramount in the care of diabetic patients.

A notable advancement in the treatment of T2DM has been the development, and increased usage, of glucagon-like peptide-1 (GLP-1) receptor agonists (RAs), due to their dual glycemia benefits and cardio-protective qualities^23^.

Multiple human clinical trials have consistently demonstrated the ability of GLP- 1RAs to lower BP^2–4,24–30^. The precise mechanisms by which GLP-1RAs elicit their BP-lowering effects remain incompletely elucidated, although the kidney, vasculature, and nervous system have shown to play significant roles^31–34^.

Furthermore, there is limited data on the effect GLP-1RAs have on nondipping BP. Gill *et al*.’s report of a trend towards diminished nighttime systolic BP, with the GLP-1RA exenatide, in T2DM subjects provides preliminary insight, but warrants further investigation^35^. To address these gaps in knowledge, we investigate the potential roles of food intake and the sympathetic nervous system in the modulation of BP’s rhythm.

Apart from its role in stimulating insulin secretion, thereby reducing blood glucose, GLP-1RAs are recognized for their ability to lower food intake through slowing gastric emptying, increasing satiety, and reducing hunger^36,37^ .

Interestingly, we and others found that experimentally limiting food availability only to the inactive phase alters BP circadian rhythm markedly in several mammalian species^38–42^, implying that meal timing profoundly influences circadian BP regulation. Moreover, our investigation has revealed that aligning food availability with the normal light-dark cycle – that is, confining food access to the active dark phase – reinstates BP and sympathetic nervous system activity rhythms in *db/db* mice, a mouse model of T2DM^43^. Thus, we hypothesize that GLP-1RAs exert their BP-lowering effects, in part, by suppressing food intake and attenuating sympathetic activity. Furthermore, the timing of GLP-1RA administration might modulate the pattern of BP dipping in T2DM.

Exenatide, a short-acting GLP-1RA with a half-life of ∼2.4 hours, was employed to test this hypothesis in *db/db* mice. These mice are a widely utilized monogenic model of T2DM, featuring an inactivation mutation in the leptin receptor, culminating in obesity, insulin resistance, and pronounced hyperglycemia^44^.

Importantly, *db/db* mice manifest disrupted circadian rhythms of both food intake and BP^43,45–48^. We administered exenatide either at the onset of the inactive light phase (zeitgeber time [ZT] 0, the light-on time) or at the start of the active dark phase (ZT12, the light-off time), with or without food. The effects on circadian BP rhythms, food intake, and urinary norepinephrine excretion were monitored. Our results indicate that, while achieving a similar reduction in BP, the timing of exenatide administration in *db/db* mice produces significant changes in BP dipping pattern. These effects are mediated through the modulation of food intake and the activity of the sympathetic nervous system. Our results suggest a role for short half-life GLP-1 RA administration timing for treating nondipping BP in diabetic patients.

## Methods

### Animals

Male BKS.Cg-Dock7^m^ +/+ Lepr^db^/J (*db/db*+/+) (Strain #: 000642) mice and age-matched male, lean nondiabetic db/+ heterozygous littermates (used as controls) were purchased from Jackson Labs (Bar Harbor, ME, USA). Mice were kept on a 12:12 light: dark cycle and fed ad libitum on a standard chow diet with free access to water unless otherwise specified. All experiments were conducted in mice between 15-19 weeks of age. All animal procedures were approved by the Institutional Animal Care and Use Committee of the University of Kentucky.

### Telemetric measurement of the BP, heart rate, and locomotor activity

Mice were implanted with a radio-telemetry transmitter (TA11PA-C10; Data Sciences International; St. Paul, MN) into the left carotid artery and allowed to recover for 10 days, as previously described^48–50^. Systolic and diastolic pressure were collected in fully conscious and free-moving *db/db* mice and age-matched nondiabetic controls (db/+).

### Food intake measurement

To continuously monitor food intake, mice were singly housed in BioDAQ cages. These cages are designed to measure food consumption and are monitored by BioDAQ software (Research Diet, New Brunswick, NJ). In addition to total food intake, the following parameters were recorded for each mouse: number of bouts, amount consumed per bout, and bout length.

For ad libitum feeding experiments, exenatide (20 µg/kg) was administered intraperitoneally at either the zeitgeber time (ZT) 0 or ZT12 (*db/db* n=8, nondiabetic control n=7). This dose and injection schedule were used for all subsequent experiments.

### Active time-restricted feeding (ATRF)

The BioDAQ system is equipped with automated gates that control access to food. These gates can be programmed to close and open at designated times. A separate cohort of mice (*db/db* n=9) was used for active time-restricted feeding (ATRF) experiments. Gates were programmed to open at ZT12 and close at ZT0, limiting food availability to only the active dark phase.

### Sympathetic nervous system measurement

In a separate cohort of mice (*db/db* n=8-9, nondiabetic control n=6), six-hour urine samples were collected across 24 hours after an acclimation period of 5 days in metabolic cages (Tecniplast, West Chester, PA) while on a gel diet (DietGel 76A; Clear H2O, Westbrook, ME) as previously described^42^. The concentration of urinary norepinephrine was measured using HPLC as previously described^51,52^ with some modifications. Briefly, urine samples were pretreated with a nonpolar cartridge according to the manufacturer’s instructions (Part # 12109301, Aligent Technologies, Santa Clara, CA). 50 µl of the eluent from the cartridge was injected and analyzed at room temperature using a 5 μm BDS Hypersil C18 (150 X 4.6 mm I.D.) column (Thermo Fisher Scientific). Mobile phase A consisted of 0.05% trifluoroacetic acid (TFA)–methanol (99.9 :0.1, v/v), and mobile phase B consisted of 0.05% TFA–methanol (40 :60, v/v). Native fluorescence was detected using excitation at 220 nm and emission at 320 nm. The total content of the urinary norepinephrine was calculated by the concentration (ng/mL) X the urine volume (mL).

In a separate cohort of mice (*db/db* n=5-6) with radio-telemetry implanted, mecamylamine (5mg/kg) or prazosin (1mg/kg) was injected at ZT0, with or without food, and with or without exenatide.

### Cosinor analysis of rhythms

The midline-estimating statistic of a rhythm (MESOR), amplitude (highest point of the rhythm from the MESOR), robustness (a measure of a continuous signal to maintain rhythmicity), and acrophase (the time at which the rhythm peaks) of rhythms for MAP and food intake were determined by cosinor analysis software (Circadian Physiology, 2^nd^ Ed., Robert Refinetti).

### Statistical analysis

All data was expressed as mean ± SEM. For comparing 1 parameter between 2 groups (i.e., ZT0 vs. ZT12), 2-tail Student’s t-test was used. For comparing 1 parameter between >2 groups (i.e., control vs. treatment groups), a one-way ANOVA was used. For comparing 2 parameters between >2 groups (i.e., light and dark phases between control vs. treatment groups), a two-way ANOVA was used. To control for family-wise error rate when comparing groups between timepoints (i.e., 2-hour MAP plots), Sidak’s multiple comparisons post hoc analysis following repeated measures ANOVA was used. For group comparisons analyzed either by one-way or two-way ANOVA, Tukey’s post hoc analysis was used. For circular data (i.e., acrophase plots) Watson’s U^2^ test was used^53^. All statistical analysis was performed by Prism 10 software (GraphPad Software; San Diego, CA), with the exception of Watson’s U^2^ test which was conducted using Oriana4, with P < 0.05 being defined as statistically significant.

### Data and Resource Availability

The data supporting this study’s findings are available from the corresponding author upon reasonable request.

## Results

### ZT0 administration of exenatide restores BP rhythmicity and dipping in *db/db* mice, while ZT12 administration worsens it

To investigate the potential effects of GLP-1-based therapy on BP rhythm in *db/db* mice, a short-acting GLP- 1 receptor agonist (GLP-1RA), exenatide (half-life ∼2.4 hours), was used. Initially, we generated dose-response curves for blood glucose and BP to exenatide to establish the appropriate dosage for subsequent experiments. The results revealed that exenatide dose-dependently reduced blood glucose (Supplemental Fig. S1A) and BP (Supplemental Fig. S1B) with similar efficacy in *db/db* mice.

These effects reached a plateau at 20 µg/kg dosage, which was used for all subsequent experiments. We then investigated the effect of time of administration on the glucose-lowering efficacy of exenatide. We injected 20 µg/kg exenatide at either the start of the light (ZT0) or dark (ZT12) phase. No difference in glucose-lowering efficacy was observed between the two-time points (Supplemental Fig. S1C). To investigate how the timing of exenatide administration influences BP’s circadian rhythm, we administered exenatide either at ZT0 or ZT12 for 3 consecutive days. Consistent with previous findings^42,43^, mean arterial pressure (MAP) displayed a robust 24-h rhythm in nondiabetic controls (Fig 1A, 1B; black), while *db/db* mice exhibited loss of rhythmicity and dipping pattern (Fig. 1A, 1B; red). Daily administration of exenatide at ZT0 restored MAP dipping (Fig. 1A, 1B; blue). In contrast, administration of exenatide at ZT12 worsened MAP dipping to reverse dipping (Fig. 1A, 1B; dark red) – a pattern characterized by BP dipping during the active/dark phase rather than the resting/light phase (Fig. 1C). However, both administration times resulted in equivalent reductions in 24-hour MAP (Fig. 1D), indicating the BP-lowering efficacy does not differ dependent upon the time of administration. The opposing effects on the BP dipping pattern of exenatide are due to the time of administration: that is, light-phase MAP decreased significantly after ZT0 exenatide administration, while dark-phase MAP decreased significantly after ZT12 exenatide administration (Fig. 1E). Similar effects were observed with systolic BP (SBP) and diastolic BP (DBP) (Supplemental Fig. S2, S3). No significant change in bodyweight was observed over the 3-day injection period: a mean loss of 0.4 and 0.2 grams after ZT0 and ZT12 exenatide administration, respectively. To verify the BP-lowering effect is attributable to exenatide, we injected the same volume of vehicle at ZT0 and ZT12. We found no observable difference in MAP after saline treatment, at either timepoint, in *db/db* mice (Supplemental Fig. S4), indicating exenatide is associated for the observed BP reduction.

**Figure 1.**
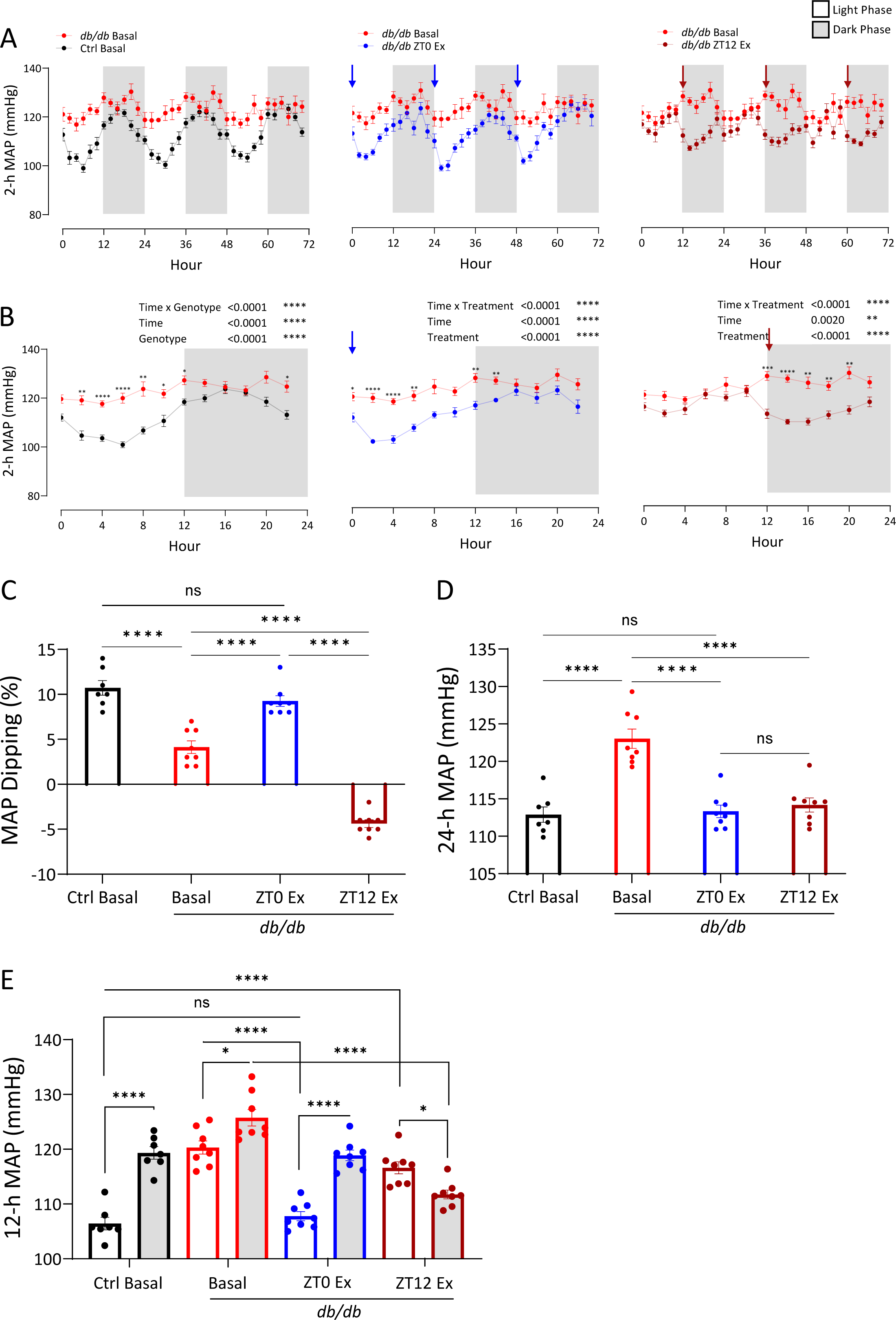
ZT0 administration of exenatide restores BP dipping in *db/db* mice while ZT12 administration worsens BP rhythm to reverse dipping. [A] 2-hour MAP bins collected over 72 hours in nondiabetic control mice (Ctrl) (black) and db/db mice under basal conditions (red), with ZT0 administration of exenatide (blue), or with ZT12 administration of exenatide (dark red). Arrows in blue or dark red indicate ZT0 or ZT12 injection of exenatide (20 µg/kg) respectively. **[B]** 2-hour MAP bins over 72-hours averaged to one 24-hour period over time. **[C]** MAP dipping calculated as [MAP (dark)-MAP (light)]/MAP (24-hour). **[D]** 72-hours of MAP averaged to one 24-hour period. **[E]** 72-hours of MAP averaged to light/dark phases (12-h). Data were analyzed by one-way ANOVA (C, D) with Tukey’s multiple comparisons post-hoc analysis, or two-way ANOVA (B, E) with Sidak’s (B) or Tukey’s (E) multiple comparisons post-hoc analysis. All data were expressed as the mean ± SEM. *P < 0.05; **P < 0.01; ***P < 0.001, ****P < 0.0001; ns, not significant.

To investigate whether the BP-lowering effect of exenatide can also be observed in normotensive nondiabetic mice, exenatide was injected at ZT0 and ZT12.

Interestingly, while ZT12 exenatide administration significantly reduced dark phase MAP, SBP, and DBP, ZT0 exenatide administration did not (Supplemental Fig. S5, S6, and S7). ZT12 exenatide administration resulted in a small, but significant reduction in 24-h MAP(Supplemental Fig. S5D). Both ZT0 and ZT12 exenatide administration lowered 24-h SBP equivalently (Supplemental Fig. S6D), with no change in 24-h DBP (Supplemental Fig. S7D).

### MAP cosinor analysis reveals improvements to circadian parameters with ZT0 but not ZT12 exenatide administration

Further quantification of the characteristics of MAP circadian rhythm by cosinor analysis revealed that exenatide administration at ZT0 significantly increased robustness (Fig. 2A) and amplitude (Fig. 2B). In contrast, exenatide administration at ZT12 had no significant effects on the MAP rhythm robustness (Fig. 2A) nor amplitude (Fig. 2B). These divergent effects on MAP robustness and amplitude are not due to different MAP-lowering effects of ZT0 vs. ZT12 exenatide administration, as both reduce the MAP MESOR to equivalent extents (Fig. 2C). Nondiabetic control acrophase did not significantly differ from *db/db* basal conditions (ZT16.6, nondiabetic control vs. ZT15.9, *db/db* basal), and ZT0 exenatide administration did not significantly alter it (ZT16.3) (Fig. 2D). Notably, variability decreased with ZT0 exenatide administration from basal conditions (SEM: 0.59, *db/db* basal vs. 0.27, ZT0 exenatide), resembling nondiabetic controls more (SEM: 0.31). ZT12 exenatide administration significantly altered acrophase, shifting it 13.8 hours forward from basal conditions, peaking during the light phase instead of the dark phase. This data shows equivalent BP-reducing efficacy from exenatide at ZT0 or ZT12 but a nearly opposite effect on BP circadian patterns based on injection time (Fig. 2E).

**Figure 2.**
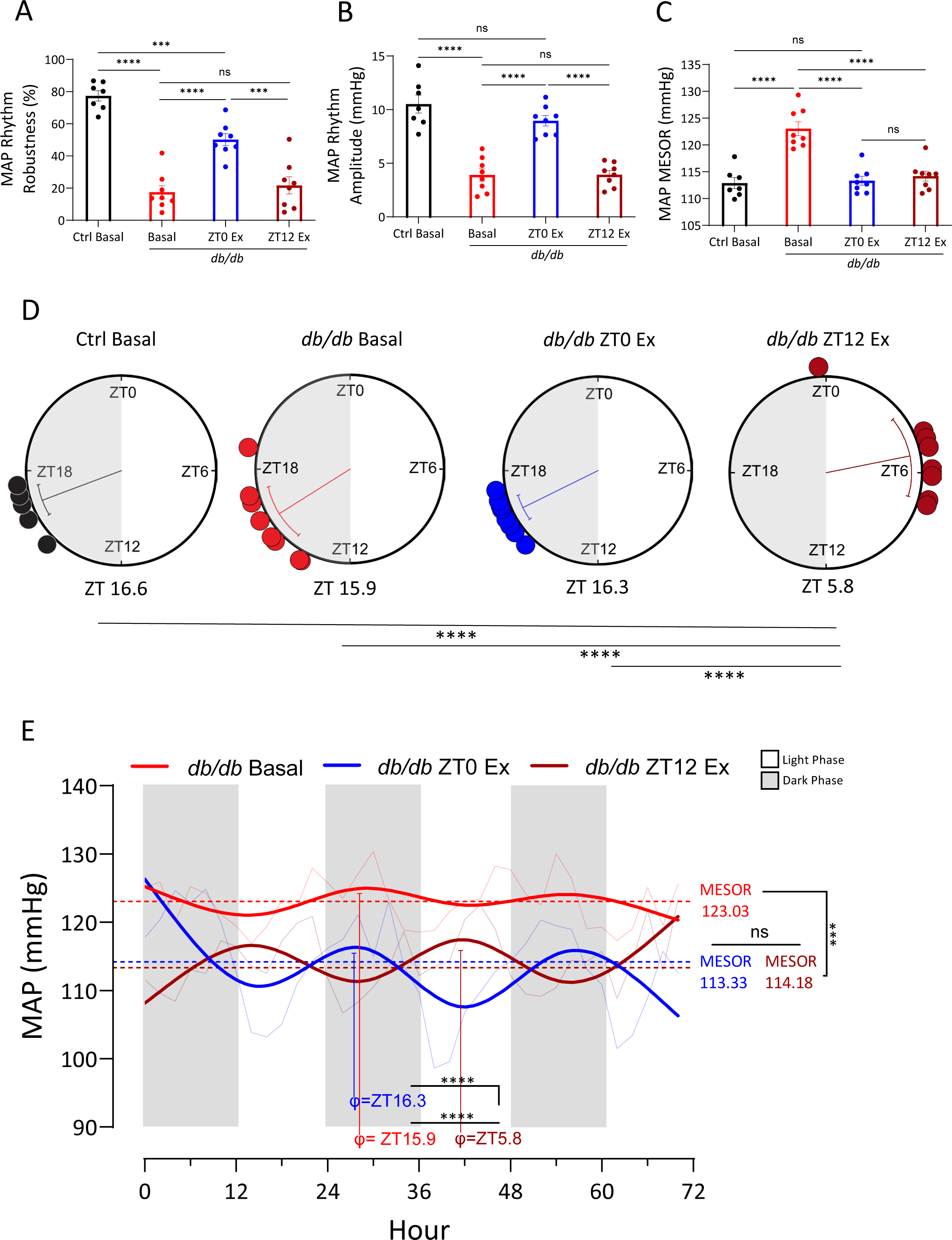
MAP cosinor analysis reveals improvements to circadian parameters with ZT0 exenatide administration. The circadian rhythm characteristics of daily MAP was examined by cosinor analysis. **[A]** MAP rhythm robustness – strength of fitted cosine curve representing rhythmic pattern of MAP. **[B]** MAP rhythm amplitude – the amount (mmHg) increase from MESOR. **[C]** MAP midline-estimating statistic of rhythm (MESOR) – the rhythm-adjusted mean. **[D]** MAP rhythm acrophase – the time (ZT) at which MAP peaks within each cycle. **[E]** Circadian plot depicting sine-fitting curves for 72-hours of *db/db* mice MAP under basal conditions (bright red), ZT0 exenatide administration (blue), and ZT12 exenatide administration (dark red), and respective MESOR and acrophase (φ). Data were analyzed by one-way ANOVA (A-C) with Tukey’s multiple comparisons post-hoc analysis. Acrophase data (D) were analyzed by Watson-Williams F- test. All data were expressed as the mean ± SEM. *P < 0.05; **P < 0.01; ***P < 0.001, ****P < 0.0001; ns, not significant.

### ZT0 administration of exenatide improves meal timing, and ZT12 administration worsens it

Previously, we reported that time-restricted feeding alters BP’s circadian rhythm^42,43^. Given the well-documented appetite-suppressive effects of endogenous GLP-1 and GLP-1RAs^54^, we sought to investigate if exenatide’s administration time differentially affected food intake patterns in *db/db* mice. Nondiabetic control mice consume 10-20% of their daily food intake during the rest/light phase (Fig. 3A, 3B; black). In contrast, *db/db* mice consume a significantly higher amount, of approximately 40%, of daily food intake during their rest/light phase (Fig. 3A, 3B; red). Administration of exenatide at ZT0 significantly suppressed food intake of *db/db* mice during the light phase (Fig. 3A, 3B; blue), whereas administration of exenatide at ZT12 significantly suppressed food intake during the dark phase (Fig. 3A, 3B; dark red). Exenatide at both administration times resulted in equivalent reductions in 24-hour food consumption (Supplemental Fig. S9A). Reduction in food intake was primarily the result of a reduced number of feeding bouts during the light or dark phase after ZT0 or ZT2 exenatide administration, respectively (Supplemental Fig. S9B and S9C). The duration of bouts did not change significantly after treatment with exenatide at either timepoint (Supplemental Fig. S9D). Cosinor analysis of food intake rhythm demonstrated that robustness (Fig. 3C) and amplitude (Fig. 3D) increased significantly with ZT0 but not with ZT12 exenatide administration.

**Figure 3.**
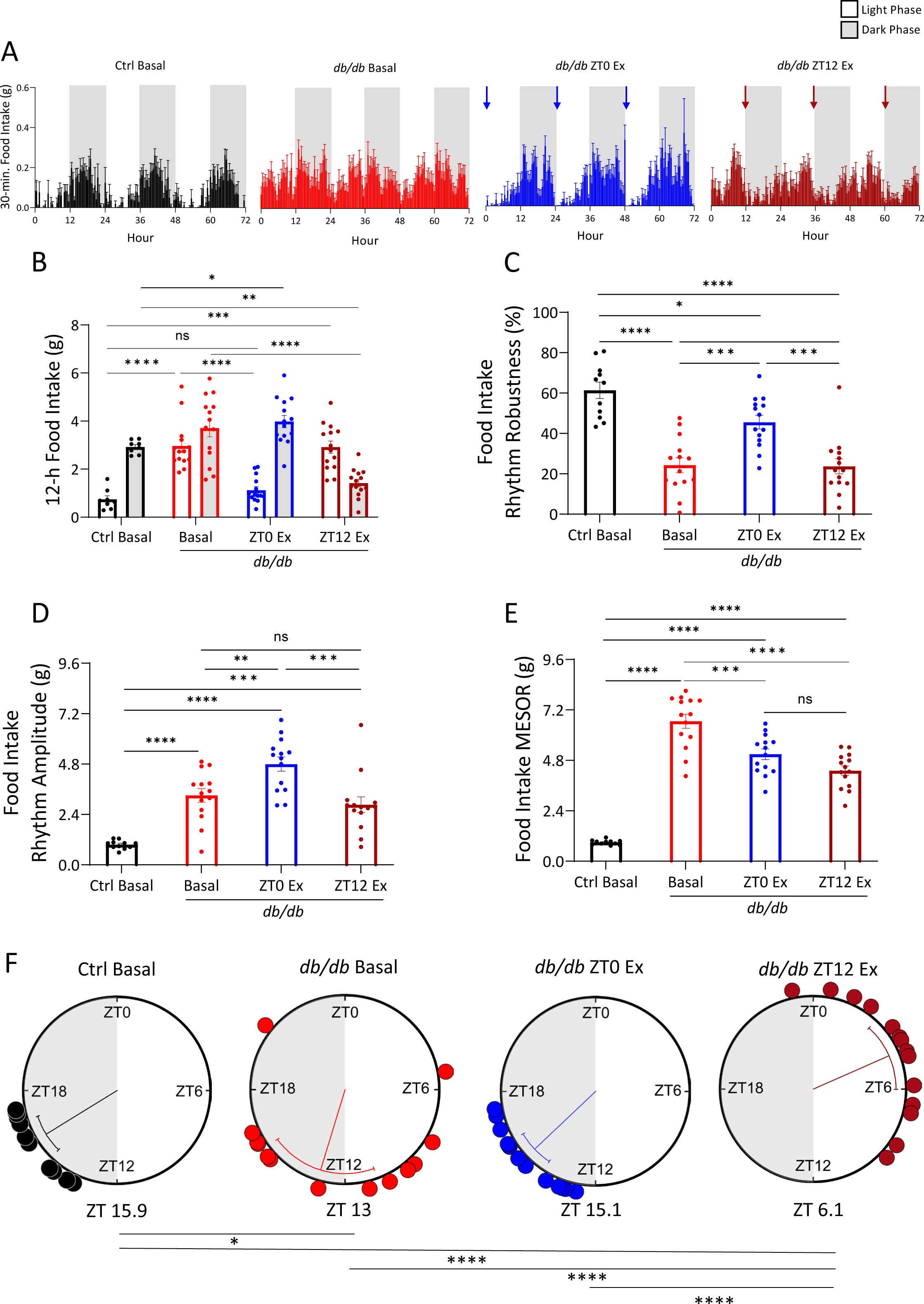
ZT0 administration of exenatide restores while ZT12 administration worsens 24-hour food intake rhythm in *db/db* mice. BioDAQ food intake monitoring system was used to measure episodic ad- libitum feeding in singly-housed nondiabetic control and *db/db* mice on a standard chow diet. **[A]** 30- minute food intake bins over 72 hours in nondiabetic control mice (Ctrl) (black), and *db/db* mice under basal conditions (red), with ZT0 exenatide (20 µg/kg) administration (blue), and with ZT12 exenatide administration (dark red). Arrows in blue or dark red indicate ZT0 or ZT12 injection of exenatide respectively. **[B]** 72-hours of food intake averaged to light/dark phases (12-h). Cosinor analysis of daily food intake are shown as **[C]** robustness, **[D]** amplitude, **[E]** MESOR, and **[F]** acrophase. Data were analyzed by two-way ANOVA (B) and one-way ANOVA (C-E) with Tukey’s multiple comparisons post-hoc analysis and were expressed as the mean ± SEM. Acrophase data (F) were analyzed by Watson-Williams F-test. *P < 0.05; **P < 0.01; ***P < 0.001, ****P < 0.0001; ns, not significant.

Reductions in food intake MESOR after ZT0 or ZT12 exenatide administration from basal conditions were equal. The acrophase of food intake rhythm significantly differed between nondiabetic control mice and *db/db* mice under basal conditions (ZT15.9, nondiabetic control vs. ZT13, *db/db* basal). ZT0 exenatide restored the food intake acrophase in *db/db* mice (ZT15.1). In contrast, ZT12 exenatide administration resulted in a significant shift forward by 16.1 hours to ZT6.1, peaking during the light phase (Fig. 3F). These changes in food intake rhythm resemble the observed changes in MAP.

In nondiabetic controls, food intake was only reduced after ZT12 exenatide administration (Supplemental Fig. S10), corresponding with the selective BP-lowering effect by ZT12 but not ZT0 exenatide administration.

### BP rhythm is temporally correlated with food intake patterns in *db/db* mice

To demonstrate the rate by which exenatide elicits its BP-lowering and food intake inhibitory effects, BP and food intake was dual-plotted in 1-minute bins starting 1 hour prior to injection at either ZT0 (Fig. 4A; blue) or ZT12 (Fig. 4B; dark red), and 3 hours after, with corresponding basal time frames. Plots depict an immediate food inhibitory effect after injection which, after a MAP spike from handling, corresponds with MAP reduction. 72-hour dual plots of MAP and food intake under basal conditions (Fig. 4C), after ZT0 (Fig. 4E), or after ZT12 (Fig. 4G) exenatide administration corroborate this, depicting similar diurnal patterns over time. Linear regression analysis demonstrates a significant correlation between MAP and food intake, regardless of the condition (Fig. 4D, 4F, and 4H). Collectively, this data suggests a potential relationship between MAP and food intake.

**Figure 4.**
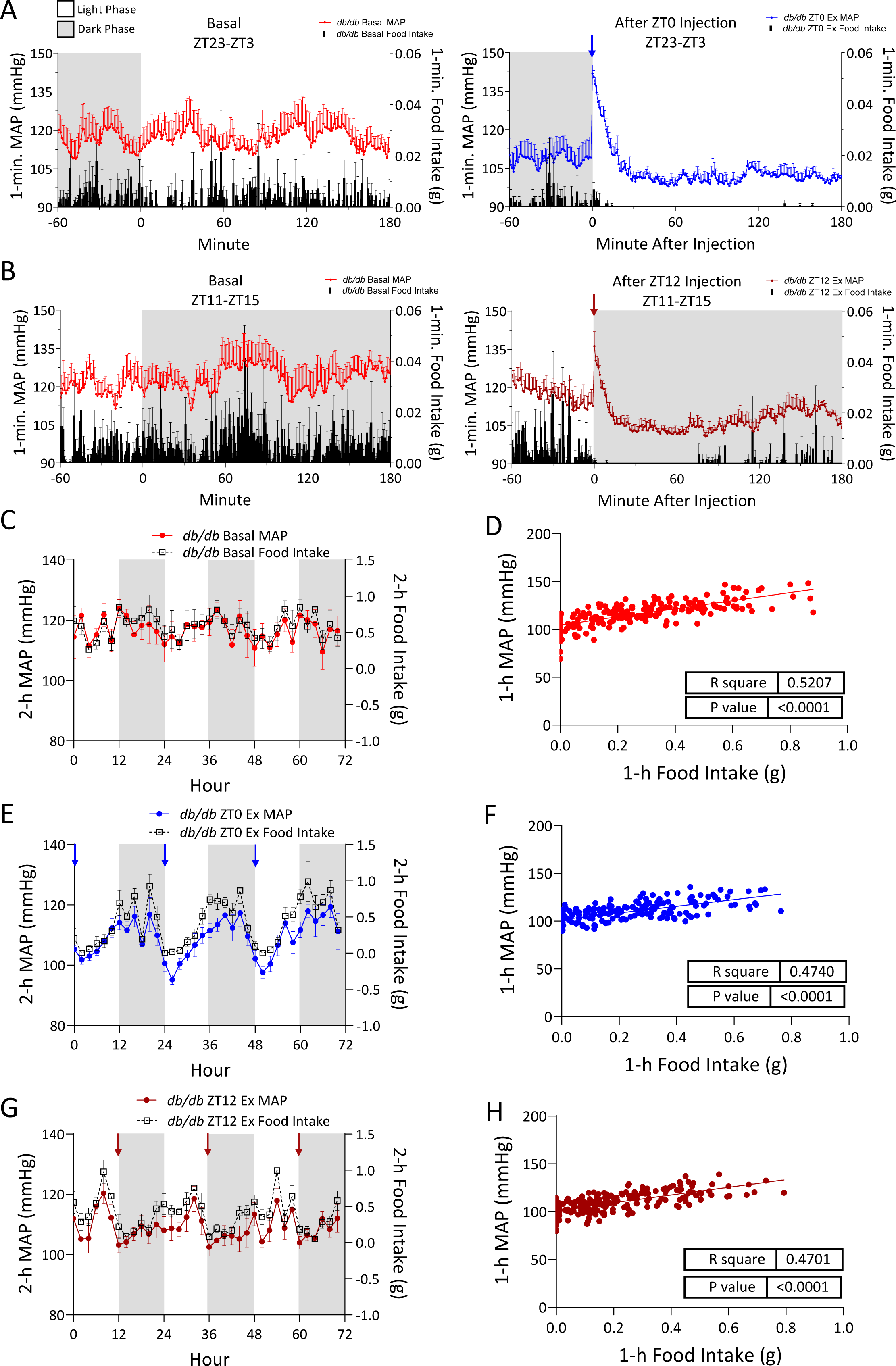
BP rhythm is temporally correlated with food intake pattern under basal conditions and with exenatide treatment in *db/db* mice. 1-minute *db/db* mean arterial pressure (MAP) and food intake bins double plotted from ZT23-ZT3 under **[A]** basal conditions (red (MAP)/black (food intake) or with ZT0 exenatide administration (blue (MAP)/black (food intake); and from ZT11-15 under **[B]** basal conditions (red (MAP)/black (food intake) or with ZT12 exenatide administration (dark red (MAP)/black (food intake). 2-h MAP and food intake bins over 72-hours were double plotted under **[C]** basal conditions, **[E]** with ZT0 exenatide administration, and **[G]** with ZT12 exenatide administration. Linear regression correlation plots of MAP and food intake under **[D]** basal conditions, **[F]** with ZT0 exenatide administration, and **[H]** with ZT12 exenatide administration.

### Exenatide loses its BP-lowering effect when food is inaccessible

We have previously reported that active time-restricted feeding (ATRF; food available only during the active/dark phase) restored BP dipping in *db/db* mice, demonstrating that food intake timing is a prominent physiological mechanism in the regulation of BP circadian rhythm^43^. Therefore, we tested if exenatide-mediated food intake reduction is responsible for its BP-lowering effect by utilizing a combined approach of ATRF and ZT0 or ZT12 exenatide administration. Similar to our previous findings, ATRF alone significantly lowered light/inactive phase MAP (Fig. 5D, days 0-5; 5E). After ZT0 exenatide injection under ATRF conditions, light phase MAP did not decrease further (Fig. 5D, days 6-9; 5E). Moreover, neither dark phase food intake (Fig. 5B, blue; 5C), number of food intake bouts (Supplemental Fig. S11B, blue; S11C), nor bout length (S11D) decreased significantly after ZT0 exenatide administration compared with ATRF alone, indicating phase-specific effects. ATRF alone did not lower dark phase MAP (Fig. 5F, days 0-5; 5G). After ZT12 exenatide administration under ATRF conditions, dark phase MAP decreased significantly (Fig. 5F, days 6-9; 5G). Dark phase food intake after ZT12 exenatide administration saw a significant reduction compared with ATRF alone (Fig. 5C), primarily as a result of reduced bout length (Supplemental Fig. 5D) but not the amount of food consumed per bout (Supplemental Fig. 5E). These results demonstrate that exenatide-induced reduction of MAP in *db/db* mice only occurs in the presence of food.

**Figure 5.**
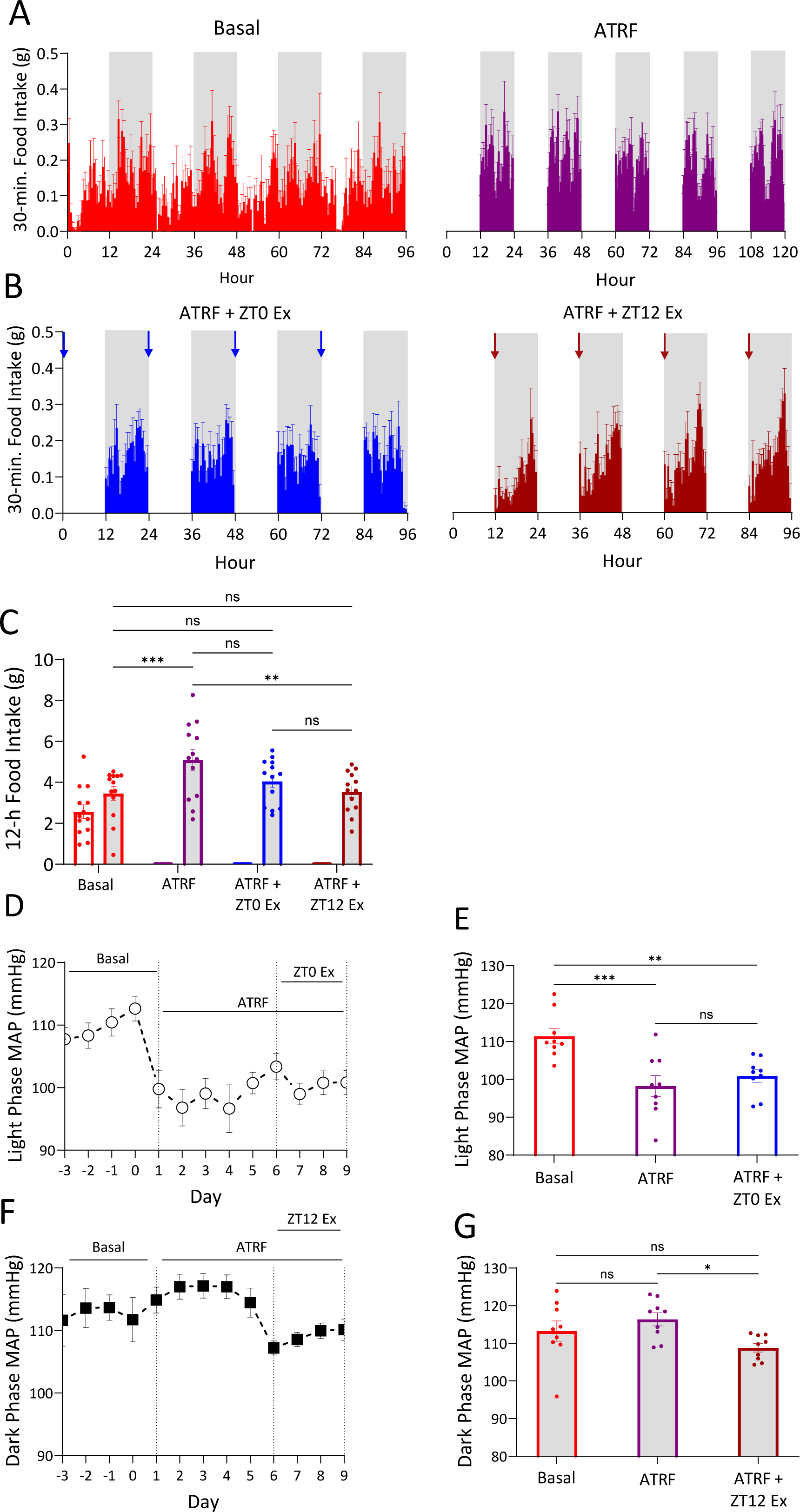
Exenatide’s BP-lowering effect is lost when food is inaccessible. BioDAQ system was programmed to limit food access to the 12-hour active dark phase only (active time-restricted feeding, ATRF). **[A]** 30- minute food intake bins in *db/db* mice over 96 and 120 hours under basal conditions (red) and under ATRF (purple), respectively. **[B]** 30-minute food intake bins over 96 hours under ATRF with ZT0 (blue) or ZT12 (dark red) exenatide administration. **[C]** Respective 30-minute food intake bins averaged to light/dark phases (12-h). **[D]** Light phase (12-h) MAP under basal (Day -3 to 0), ATRF only (Day 1 to 5), and ATRF with ZT0 exenatide administration (Day 6 to 9). **[E]** Respective light phase MAP days averaged. **[F]** Dark phase (12-h) MAP under basal (Day -3 to 0), ATRF only (Day 1 to 5), and ATRF with ZT12 exenatide administration (Day 6 to 9). **[G]** Respective dark phase MAP days averaged. Data were analyzed by two-way ANOVA (C) and one-way ANOVA (E,G) with Tukey’s multiple comparisons post-hoc analysis and were expressed as the mean ± SEM. *P < 0.05; **P < 0.01; ***P < 0.001, ****P < 0.0001; ns, not significant.

**Figure 6.**
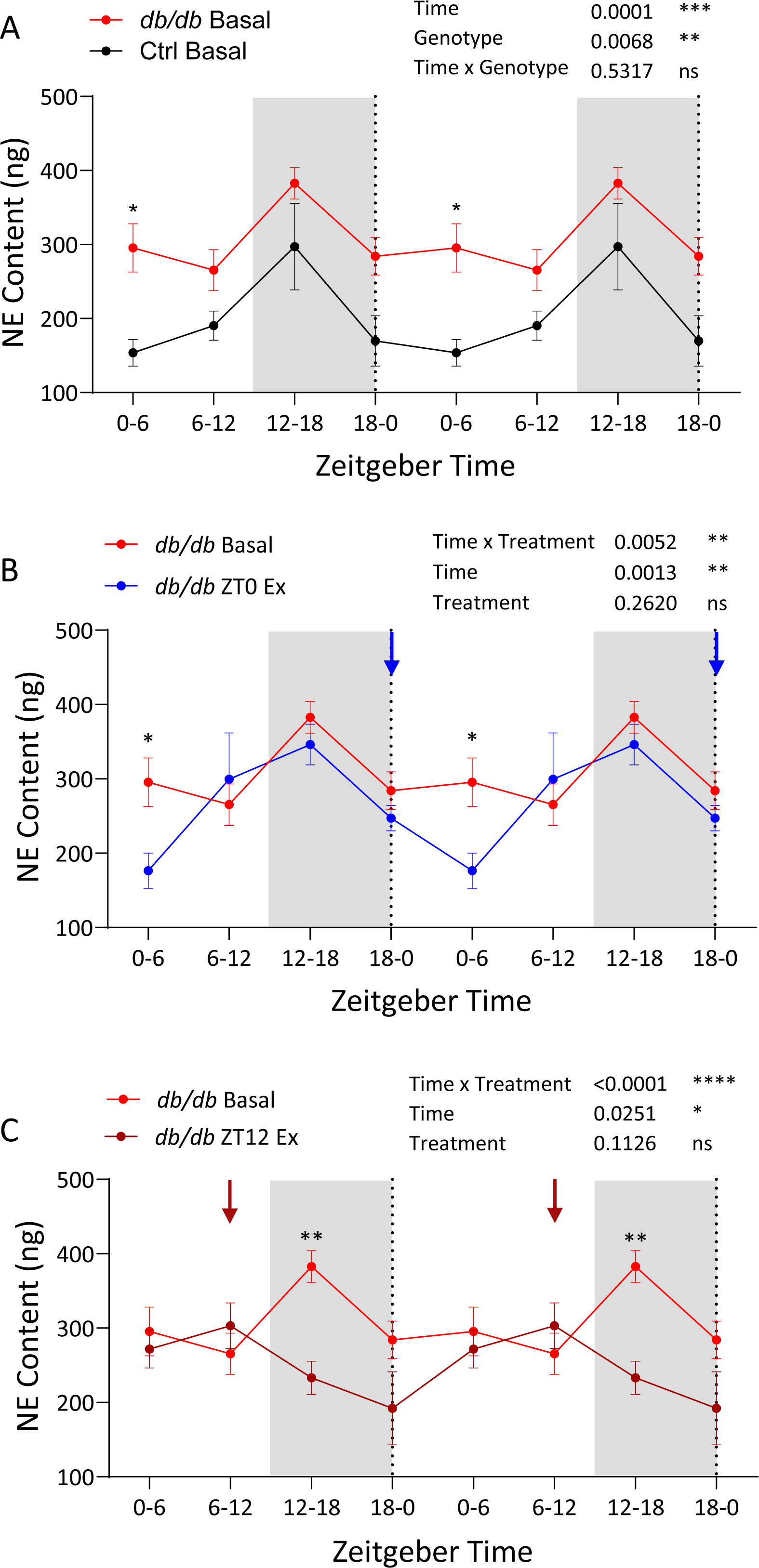
Exenatide has sympathetic nervous system-suppressing properties, with ZT0 administration improving, and ZT12 exacerbating, urinary norepinephrine (NE) excretion diurnal rhythm in *db/db* mice. Urinary NE content of **[A]** nondiabetic control mice (Ctrl) and *db/db* mice under basal conditions; [B] *db/db* basal and after ZT0 exenatide. [C] *db/db* basal and after ZT12 exenatide administration. Levels of NE are expressed as total content, calculated by concentration (ng/mL) x urine volume (mL). Graphs were double- plotted to better depict the daily changes in NE content over time. Arrows indicate the time of exenatide injection. Data were analyzed by two-way ANOVA with Sidak’s (B,C,D) multiple comparisons post-hoc analysis and were expressed as the mean ± SEM. *P < 0.05; **P < 0.01; ***P < 0.001; ns, not significant.

### Exenatide-induced BP reduction is associated with reduced sympathetic nervous system (SNS) activity

Our previous report demonstrated that food intake inhibition lowers sympathetic nervous system (SNS) activity, thus reducing BP in *db/db* mice^43^. We, therefore, explored whether exenatide-induced reductions in food intake and BP were associated with the suppression of SNS by measuring urinary norepinephrine (NE) excretion; as well as administering exenatide after ganglionic blockade with mecamylamine or alpha-1 adrenergic antagonism with prazosin, in the presence or absence of food. Urinary NE excretion was measured in 6-hour urine samples across 24 hours at baseline, and after exenatide injection at ZT0 or ZT12 in nondiabetic control and *db/db* mice. Under basal conditions, *db/db* mice have significantly higher urinary NE content than nondiabetic controls (Fig. 6A). Exenatide, whether administered at ZT0 or ZT12, had a significant suppressive effect on urine NE content for the 6 hours following injection (Fig. 6B; 6C). Exenatide-induced NE excretion suppression occurred mostly within 6 hours post injection. As such, ZT0 administration improved light-dark phase differences, whereas ZT12 administration worsened NE excretion rhythm in *db/db* mice. Interestingly, exenatide treatment in nondiabetic control mice resulted in a delayed suppressive effect after ZT0 administration, and a blunting of rhythm after ZT12 administration.

Mecamylamine, a nicotinic receptor antagonist, was injected at ZT0 and resulted in significant reduction in MAP from basal conditions in *db/db* mice (Fig. 7B; 7D).

**Figure 7.**
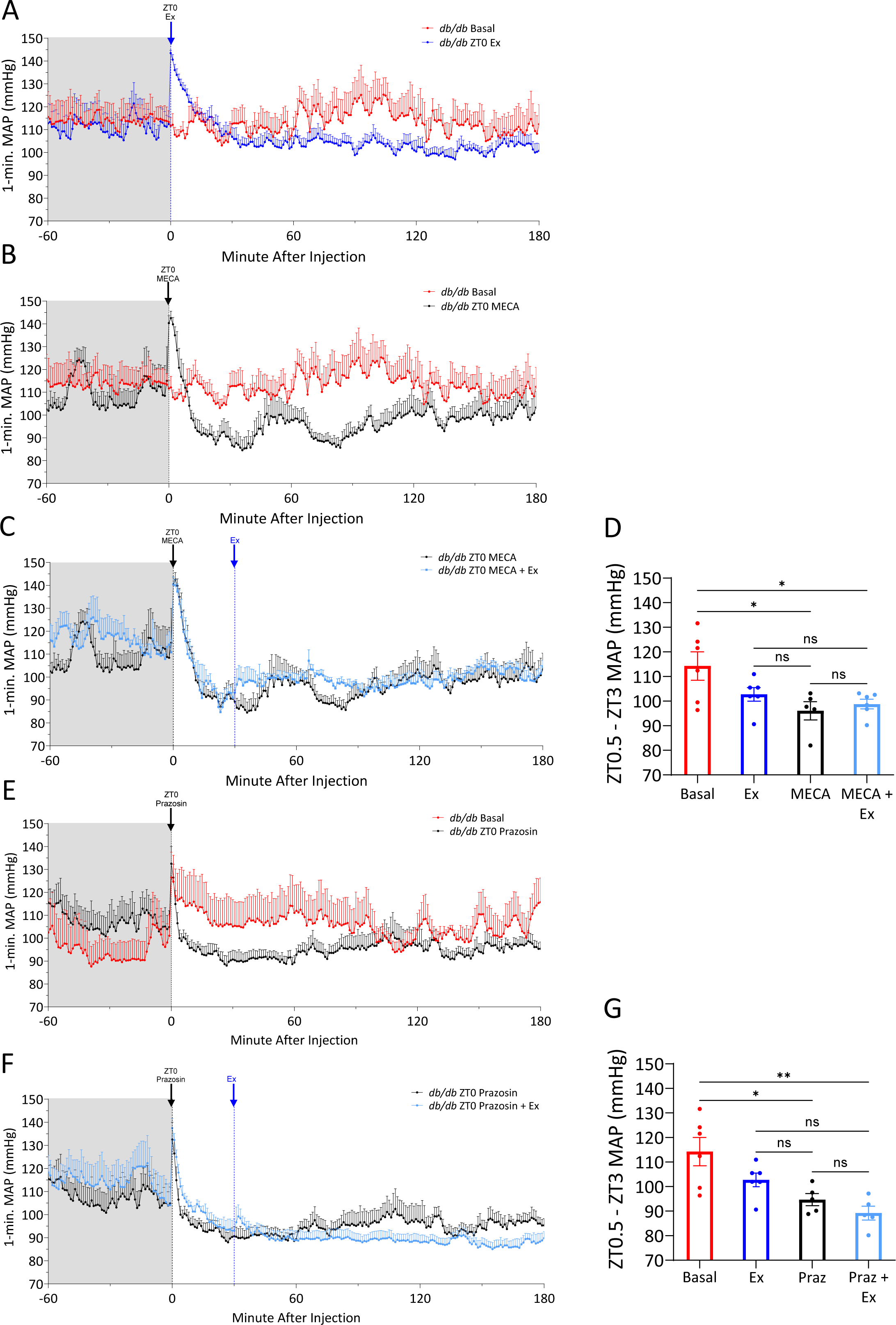
Under ad libitum feeding, the addition of exenatide does not result in further MAP decrease after ganglionic blockade with mecamylamine, and is greatly attenuated after alpha-1 adrenergic antagonism with prazosin in *db/db* mice. 1-minute *db/db* MAP bins of **[A]** basal (red) and ZT0 exenatide (20 µg/kg) administration (blue), **[B]** basal (red) and ZT0 mecamylamine administration (MECA, 5mg/kg, black). **[C]** ZT0 MECA (black) administration and ZT0 MECA with exenatide (light blue) administration. **[D]** Respective MAP recordings averaged from ZT0.5 to ZT3. ZT0.5 chosen to exclude stress-induced MAP spike; ZT3 chosen as cutoff as MECA’s MAP-lowering effect’s last approximately 3 hours. 1-minute *db/db* MAP bins of **[E]** basal (red) and ZT0 of prazosin (1mg/kg, black) administration, **[F]** ZT0 prazosin (black) administration and ZT0 prazosin with exenatide (light blue) administration. **[G]** Respective MAP recordings. Data were analyzed by one-way ANOVA (D,G) with Tukey’s multiple comparisons post-hoc analysis. All data were expressed as the mean ± SEM. *P < 0.05; **P < 0.01; ***P < 0.001, ****P < 0.0001; ns, not significant.

After MAP reached its lowest point, exenatide was subsequently injected. The addition of exenatide did not result in further MAP decrease (Fig. 7C; 7D).

Similarly, ZT0 administration of prazosin decreased MAP from basal conditions (Fig. 7E; 7G). When combined, the MAP-lowering effects of exenatide were greatly attenuated (Fig. 7F), resulting in nonsignificant reduction in MAP (Fig. 7G) compared with prazosin alone. These results demonstrate that exenatide loses its BP-lowering effects after pharmacologically blocking the effects of the SNS, suggesting that exenatide works through this pathway. To further build upon the relationship between exenatide’s BP-lowering effects, food intake inhibition, and SNS activity suppression, the above experimental conditions were employed after rescue of BP dipping via ATRF. Neither ZT0 administration of mecamylamine (Fig. 8A), prazosin (Fig. 8D), nor combination with exenatide (Fig. 8B; 8E) lowered MAP further during the fasting phase than ATRF alone (Fig. 8C; 8F). Thus, when food is inaccessible, SNS-mediated BP regulation is attenuated – demonstrated by the lack of MAP reduction after pharmacological blocking of the SNS. As such, the addition of exenatide does not lower BP under these conditions.

**Figure 8.**
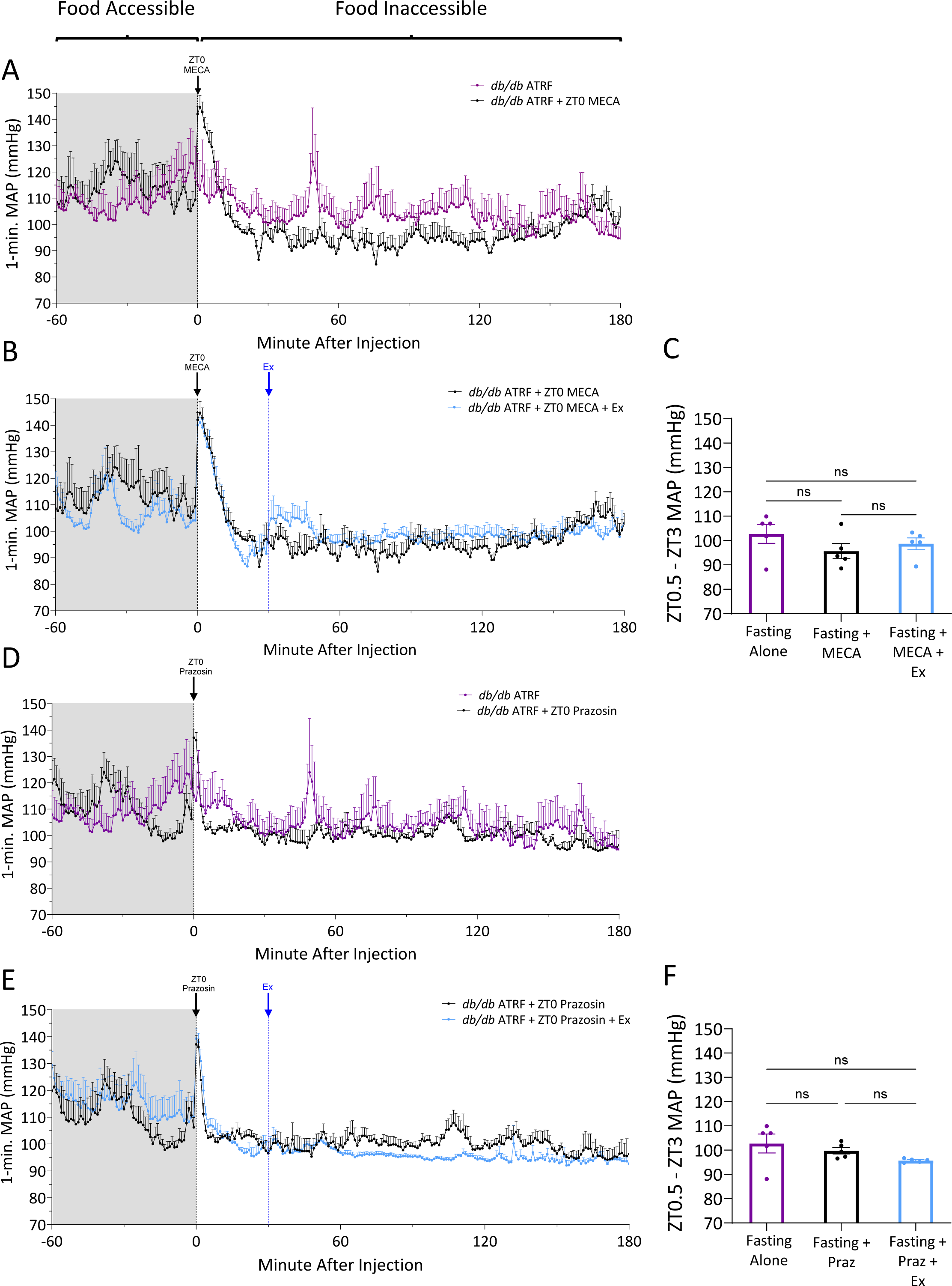
Neither ZT0 mecamylamine nor prazosin, with or without exenatide, reduces BP further under ATRF conditions during the light phase/fasting period in *db/db* mice. BioDAQ system was programmed to limit food access to the 12-hour active dark phase only (active time-restricted feeding, ATRF). 1-minute *db/db* MAP bins of **[A]** ATRF (purple) and ATRF with ZT0 mecamylamine (MECA, 5mg/kg, black) administration, **[B]** ATRF with ZT0 MECA (black) administration and ATRF with ZT0 MECA plus exenatide (light blue) administration. **[C]** Respective MAP recordings averaged from ZT0.5 to ZT3. ZT0.5 chosen to exclude stress-induced MAP spike; ZT3 chosen as a cutoff as MECA’s MAP-lowering effect’s last approximately 3 hours. 1-minute *db/db* MAP bins of **[D]** ATRF (purple) and ATRF with ZT0 prazosin (1mg/kg, black) administration, **[E]** ATRF with ZT0 prazosin (black) and ATRF with ZT0 prazosin plus exenatide (light blue) administration. **[F]** Respective MAP recordings averaged from ZT0.5 to ZT3. Data were analyzed by one-way ANOVA (C,F) with Tukey’s multiple comparisons post-hoc analysis. All data were expressed as the mean ± SEM. *P < 0.05; **P < 0.01; ***P < 0.001, ****P < 0.0001; ns, not significant.

## Discussion

The high prevalence of hypertension and nondipping BP contribute to cardiovascular-related death in T2DM^8,12,13,55^. GLP-1RA-based therapy not only lowers blood glucose but BP as well^56^. Here, we sought to determine the effects of exenatide, a short half-life GLP-1RA, on BP circadian rhythm when administered at either the onset of the rest/light phase (ZT0) or dark/active phase (ZT12). As a model of T2DM and nondipping BP, diabetic *db/db* mice were subjected to a series of ZT0 or ZT12 exenatide injections. We demonstrate that BP dipping is restored after ZT0 exenatide administration and worsened to reverse dipping when given at ZT12. Coinciding with this, *db/db* hyperphagia is reduced, and its day-to-night rhythm is restored with ZT0 exenatide administration and worsened with ZT12 administration. Given this correlation, we investigated if the changes in food intake patterns served as a potential physiological mechanism for the changes in BP. To test this, ATRF (food only accessible during the dark/active phase) was coupled with ZT0 exenatide administration. Under ATRF conditions, light phase BP-lowering was completely abolished after ZT0 exenatide administration, but still present after ZT12 exenatide administration during the dark phase. These results indicate that without food present, exenatide cannot elicit its BP-lowering effects. Furthermore, administration of exenatide at either timepoint, reduces SNS activity, demonstrated by suppression of NE content. Exenatide-induced BP-lowering was abolished following pharmacological blockade of SNS activity, with or without food. There is speculation that time-restricted eating and chronotherapy may be efficacious in the treatment of cardio-metabolic disorders. For example, Panda *et al*. found diverse therapeutic benefits for metabolic conditions associated with T2DM by limiting food availability to the first 8-9 hours of the active/dark phase in mice fed a combination of high-fat and/or high-sugar diets^57^. Several BP medications have been shown to be more effective when taken before bed compared with upon awakening^58–60^. Our results suggest that ZT0 exenatide administration is superior to ZT12 for the correction of nondipping BP in T2DM.

The popularity of GLP-1RA usage has prompted investigations into this drug class’ effects outside of glycemic control. However, the mechanisms by which GLP-1RA lowers BP are not fully understood. Activation of the GLP-1 receptor in the kidney produces a natriuretic and diuretic effect via reducing Na^+^/H^+^ exchanger 3 activity^31^ and increasing renal blood flow^32^. In the heart, GLP-1 receptor activation promotes the secretion of atrial natriuretic peptide^33^. GLP-1 RAs modulate the autonomic nervous system that transiently increases heart rate^34,35^ but decreases systolic BP^35^. One identified mechanism involves the lessening of carotid-body overstimulation witnessed in hyperglycemic states^61^. Our findings contribute to this body of work by implicating food intake inhibition as an additional mechanism. Lifestyle and pharmacological approaches to target sympathetic nervous system over-activity have been efficacious in the treatment of T2DM. It is well established that inhibiting food intake modulates the autonomic nervous system in humans and rodents^43,62^. Specifically, fasting has shown to lower catecholamine secretion, thereby suppressing SNS activity^43,63,64^. Interestingly, central administration of exenatide has been shown to increase MAP and trigger SNS activation in rats’ dose-dependently^65^. However, after exenatide treatment, we observed a reduction in urinary NE excretion and loss of BP-reducing effect after pharmacological blocking of SNS activity. These opposing effects on SNS and BP remain uncertain, but may relate to differing effects in the central nervous system versus the periphery.

Some limitations in the present study warrant further investigation for future studies. While we demonstrate herein that ZT0 exenatide administration restores BP dipping, we do not know if long-term ZT0 treatment protects against the target organ damage associated with nondipping BP^66^. Because our study utilized only males, whether exenatide has similar effects in females remains to be explored. We have previously shown that time-restricted feeding in *db/db* mice restores clock gene oscillation in tissues critical for BP regulation^43^.

Because GLP-1RAs suppress food intake, whether different exenatide timing induces changes to clock gene expression remains to be seen. Continued growth of the GLP-1RA market is expected, with longer-acting versions dominating^67^.

Our findings would suggest that, in the context of BP rhythm, there may be consequences to chronic GLP-1 receptor activation in the treatment of T2DM for those exhibiting nondipping BP. Future research into the effects of longer-acting GLP-1RA on BP circadian rhythm is necessary.

In summary, we have demonstrated that exenatide administration at ZT0 restores BP dipping in nondipping *db/db* mice, while administration at ZT12 worsens nondipping BP to reverse BP dipping. Inhibition of food intake and sympathetic activity mediate the BP-lowering effect of exenatide. Our results suggest that time of administration be a consideration in those who exhibit nondipping BP with T2DM.

## Acknowledgments

### Author Contributions

ANC performed the experiments, analyzed data, and wrote the manuscript with inputs from MG and ZG. WS performed all surgeries and contributed to data acquisition. TH participated in experiments and data analysis. MCG and ZG co- developed the hypothesis, conceived experimental ideas, contributed to the interpretation of results and discussion, critically reviewed and refined the manuscript.

### Sources of Funding

This work is supported by NIH HL141103 and HL164398 to MG and ZG. ANC is a recipient of NIH F31HL162463.

## Abbreviations & Definitions

ATRF: active-time restricted feeding; food inaccessible during light phase
BP: blood pressure
DBP: diastolic blood pressure
Ex: exenatide
GLP-1: glucagon like peptide-1
MA: mean arterial pressure
MECA: mecamylamine; non-selective, non-competitive antagonist of nicotinic acetylcholine receptors
NE: norepinephrine
Praz: prazosin; alpha-1 adrenergic receptor antagonist
RA: receptor agonist
SNS: sympathetic nervous system
SBP: systolic blood pressure
T2DM: type 2 diabetes mellitus
ZT: zeitgeber time; time scale referenced to external cues that synchronize circadian rhythms
ZT0: zeitgeber time 0; lights on
ZT12: zeitgeber time 12; lights off

