## supplemental figures for "Exenatide administration time determines the effects on blood pressure dipping in *db/db* mice via modulation of food intake and sympathetic activity"

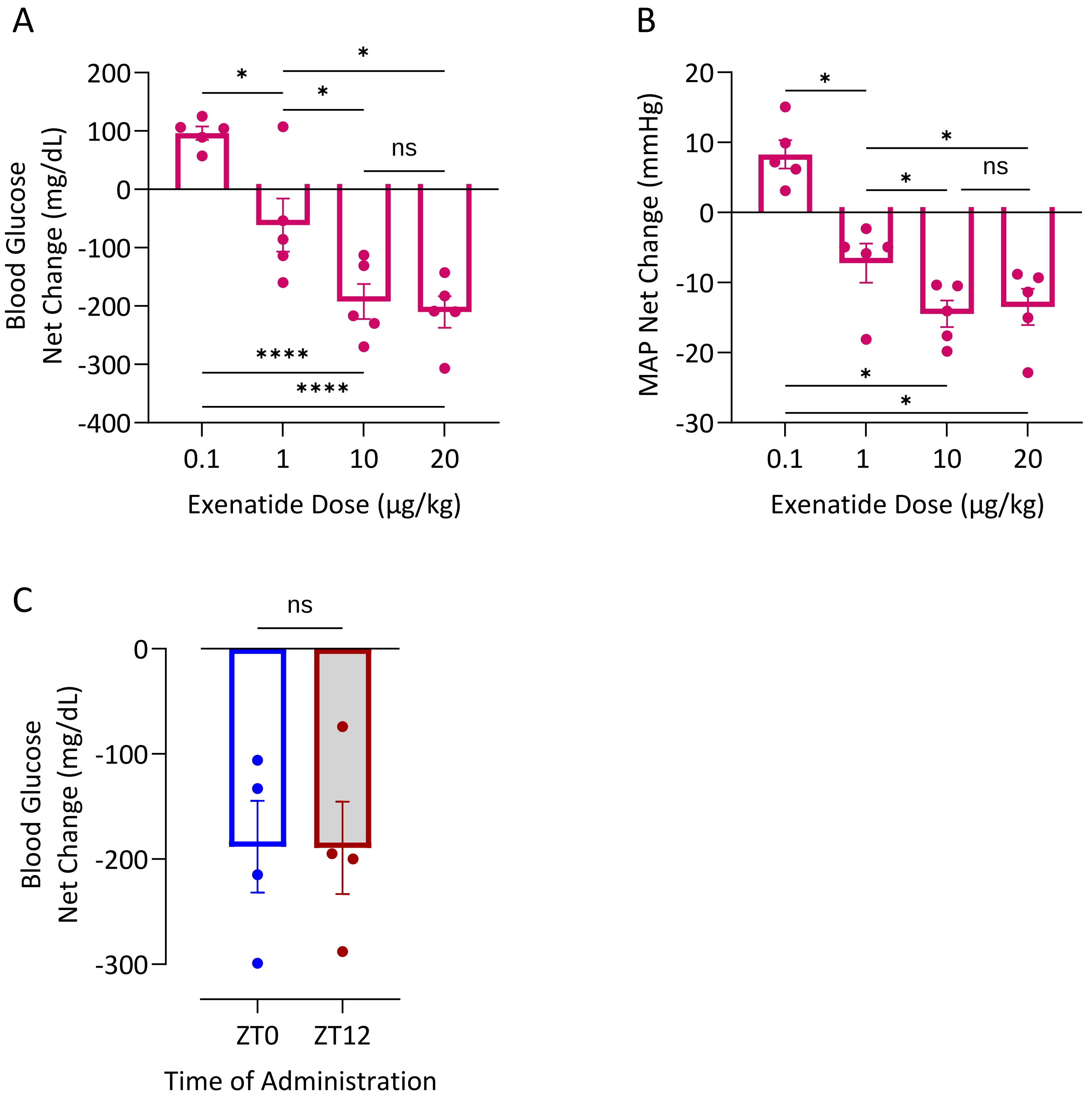


**Supplemental Figure S1. [A]** Difference in baseline blood glucose measured (via tail cut) at ZT6 and 2 hours after exenatide (0.1, 1, 10, and 20µg/kg) injection, at ZT8, in *db/db* mice. **[B]** Baseline MAP measured at ZT6 and 2 hours after exenatide (0.1, 1, 10, and 20µg/kg) injection, at ZT8, in *db/db* mice. **[C]** Difference in baseline blood glucose measured at ZT0 or ZT12 and 2 hours after exenatide (20µg/kg) injection in *db/db* mice. Data were expressed as mean ± SEM and analyzed by one-way ANOVA with Tukey's multiple comparisons post-hoc analysis (A,B) or two-tailed T test (C).*P < 0.05; **P < 0.01; ***P < 0.001, ****P < 0.0001; ns, not significant.

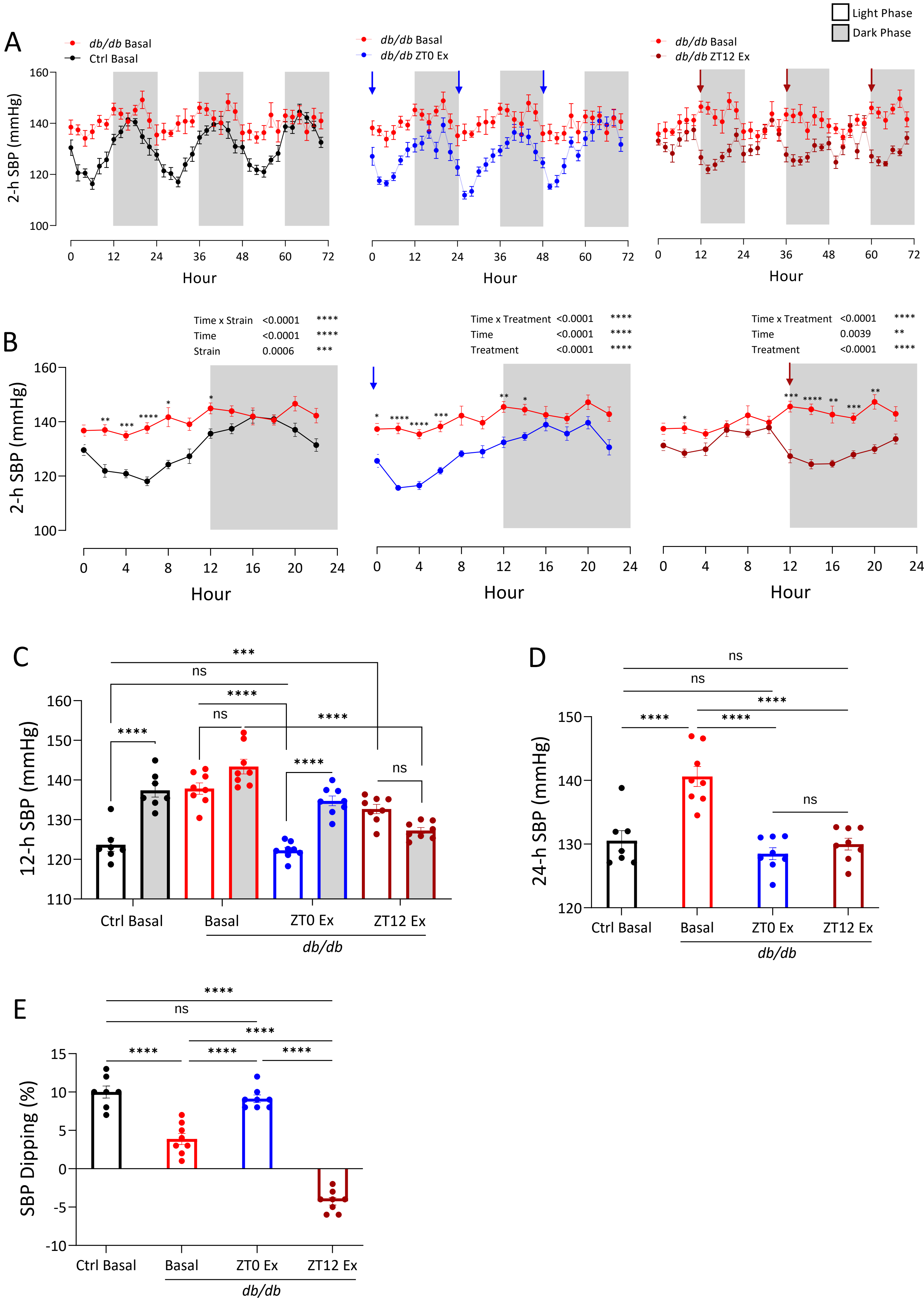


**Supplemental Figure S2. ZT0 administration of exenatide restores systolic BP (SBP) dipping in *db/db* mice while ZT12 administration worsens SBP rhythm to reverse dipping.** SBP was monitored via radiotelemetry in diabetic *db/db* and control mice. **[A]** 2-hour SBP bins collected over 72 hours in control mice basal (black), *db/db* mice basal (red), *db/db* mice with ZT0 exenatide (blue), or *db/db* mice with ZT12 exenatide (dark red). Arrows in blue or dark red indicate ZT0 or ZT12 injection of exenatide (20 μg/kg), respectively. **[B]** 2-hour SBP bins over 72-hours averaged to one 24-hour period. Arrows in blue or dark red indicate ZT0 or ZT12 injection of exenatide (20 μg/kg), respectively. **[C]** 72-hours of SBP averaged to light/dark phases (12-h). **[D]** 72 hours of SBP averaged to 24 hours. **[E]** SBP dipping calculated as [SBP (dark)-SBP (light)]/SBP(dark). Data were expressed as mean ± SEM and analyzed by two-way ANOVA with Sidak's (B) or Tukey's (C) multiple comparisons post-hoc analysis, and one-way ANOVA (D, E) with Tukey's multiple comparisons post-hoc analysis. *P < 0.05; **P < 0.01; ***P < 0.001, ****P < 0.0001; Data not indicated, or labeled as ns, is not significant.

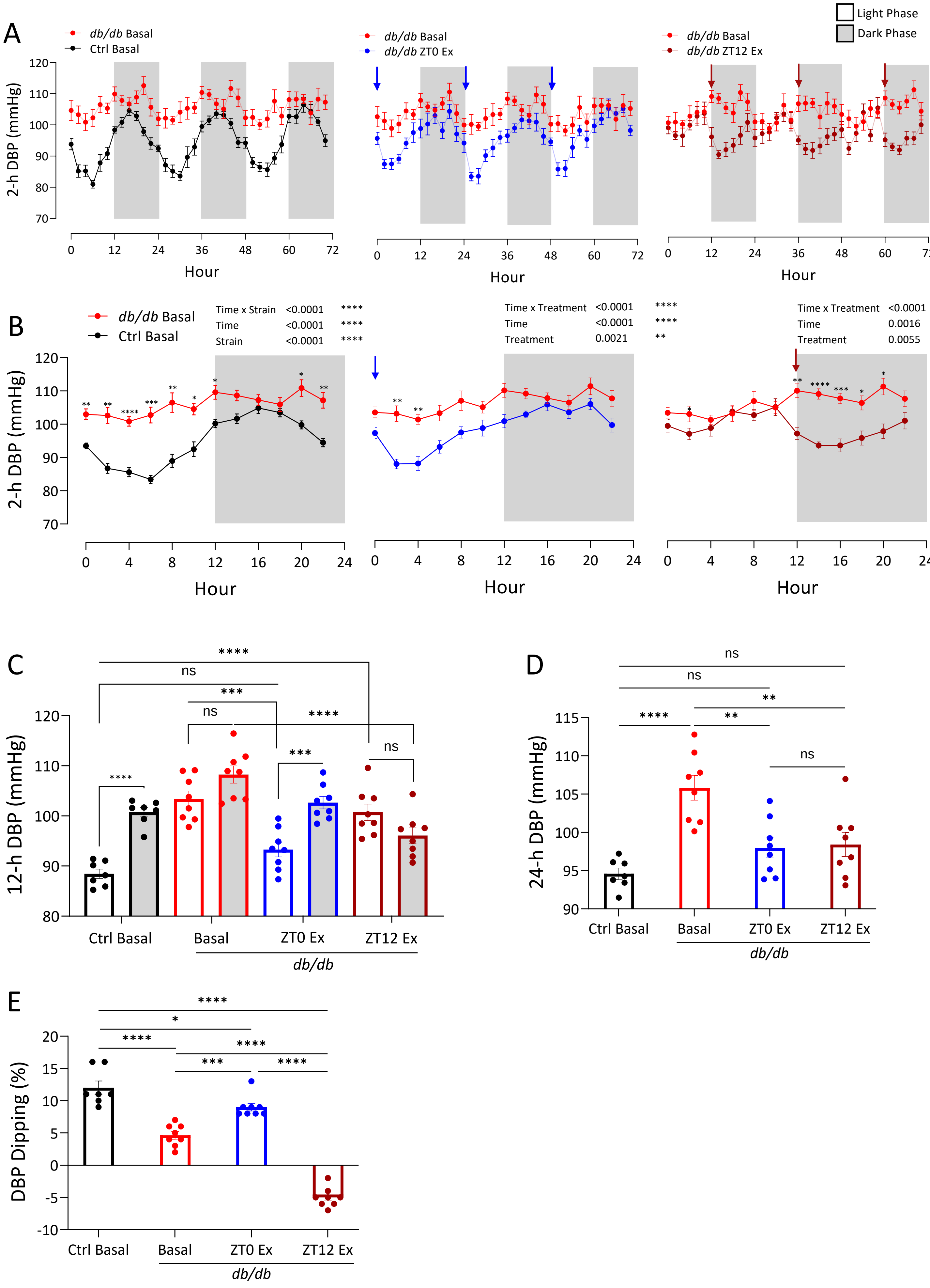


**Supplemental Figure S3. ZT0 administration of exenatide restores diastolic BP (DBP) dipping in *db/db* mice while ZT12 administration worsens DBP rhythm to reverse dipping.** DBP was monitored via radiotelemetry in diabetic *db/db* and control mice. **[A]** 2-hour DBP bins collected over 72 hours in control mice basal (black), *db/db* mice basal (red), *db/db* mice with ZT0 exenatide (blue), or *db/db* mice with ZT12 exenatide (dark red). Arrows in blue or dark red indicate ZT0 or ZT12 injection of exenatide (20 µg/kg), respectively. **[B]** 2-hour DBP bins over 72-hours averaged to one 24-hour period. Arrows in blue or dark red indicate ZT0 or ZT12 injection of exenatide (20 µg/kg), respectively. **[C]** 72-hours of DBP averaged to light/dark phases (12-h). **[D]** 72 hours of DBP averaged to 24 hours. **[E]** DBP dipping calculated as [DBP (dark)-DBP (light)]/DBP (dark). Data were expressed as mean ± SEM and analyzed by two-way ANOVA with Sidak's (B) or Tukey's (C) multiple comparisons post-hoc analysis, and one-way ANOVA (D, E) with Tukey's multiple comparisons post-hoc analysis. *P < 0.05; **P < 0.01; ***P < 0.001, ****P < 0.0001; Data not indicated, or labeled as ns, is not significant.

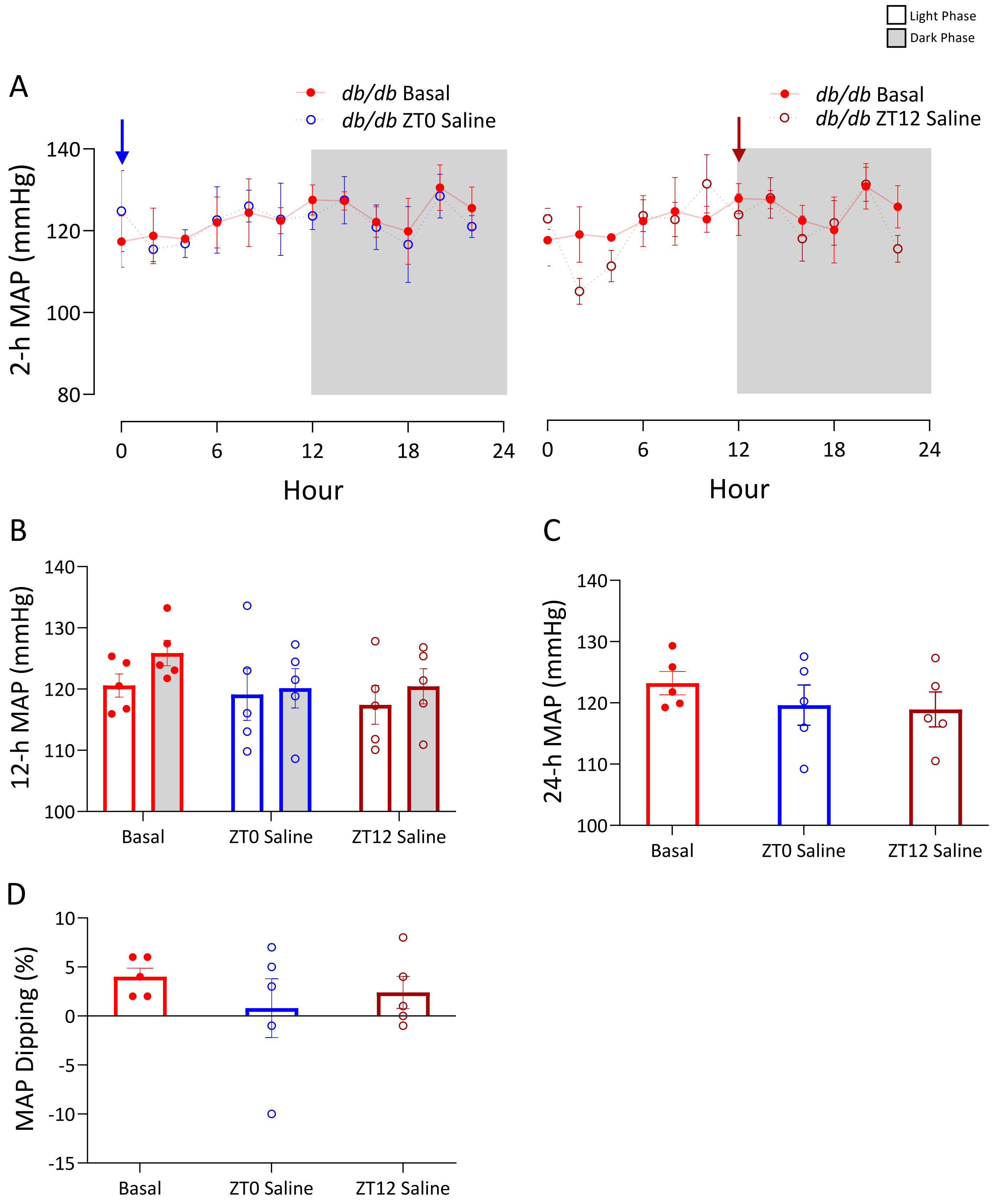


**Supplemental Figure S4**. **Neither ZT0 nor ZT12 saline administration reduces BP in *db/db* mice. [A]** 2-hour MAP bins collected over 24 hours under basal (red), after ZT0 exenatide administration (blue), or ZT12 exenatide administration (dark red). Arrows in blue or dark red indicate ZT0 or ZT12 injection of exenatide (20 μg/kg), respectively. **[B]** 12-hour bins of MAP **[C]** 24-hour average of MAP **[D]** MAP dipping - the percentage of MAP decrease from dark to light phase. Data were analyzed by two-way ANOVA with Sidak's (A) or Tukey's (B) multiple comparisons post-hoc analysis, and one-way ANOVA (C,D) with Tukey's multiple comparisons post-hoc analysis and were expressed as the mean ± SEM. *P < 0.05; **P < 0.01; ***P < 0.001, ****P < 0.0001; Data not indicated, or labeled as ns, is not significant..

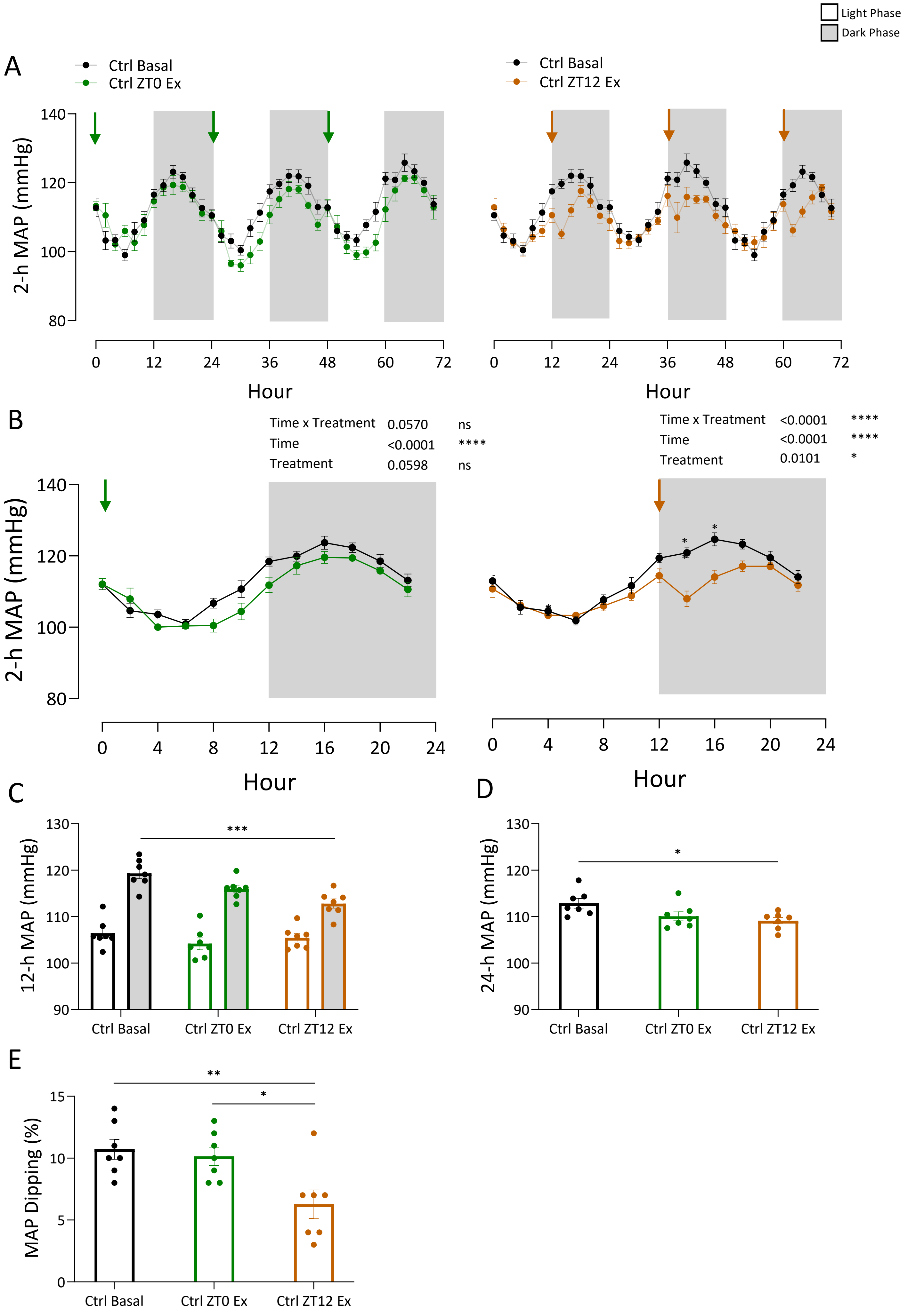


**Supplemental Figure S5. ZT12, but not ZT0, exenatide significantly decreases MAP in nondiabetic controls. [A]** 2-hour MAP bins collected over 72 hours under basal (black), after ZT0 exenatide administration (green), or ZT12 exenatide administration (orange). Arrows in green or orange indicate ZT0 or ZT12 injection of exenatide (20 µg/kg), respectively. **[B]** 2-hour MAP bins over 72-hours averaged to one 24-hour period. **[C]** 72 hours of MAP averaged to light/dark phases (12-h). **[D]** 72 hours of MAP averaged to 24 hours. **[E]** MAP dipping - the percentage of MBP decrease from dark to light phase. Data were analyzed by two-way ANOVA (B, C) or one-way ANOVA (D, E) with multiple comparisons test and were expressed as the mean ± SEM. *P < 0.05; **P < 0.01; ***P < 0.001, ****P < 0.0001; ns, not significant. Data were analyzed by two-way ANOVA with Sidak's (B) or Tukey's (C) multiple comparisons post-hoc analysis, and one-way ANOVA (D,E) with Tukey's multiple comparisons post-hoc analysis and were expressed as the mean ± SEM. *P < 0.05; **P < 0.01; ***P < 0.001, ****P < 0.0001; Data not indicated, or labeled as ns, is not significant.

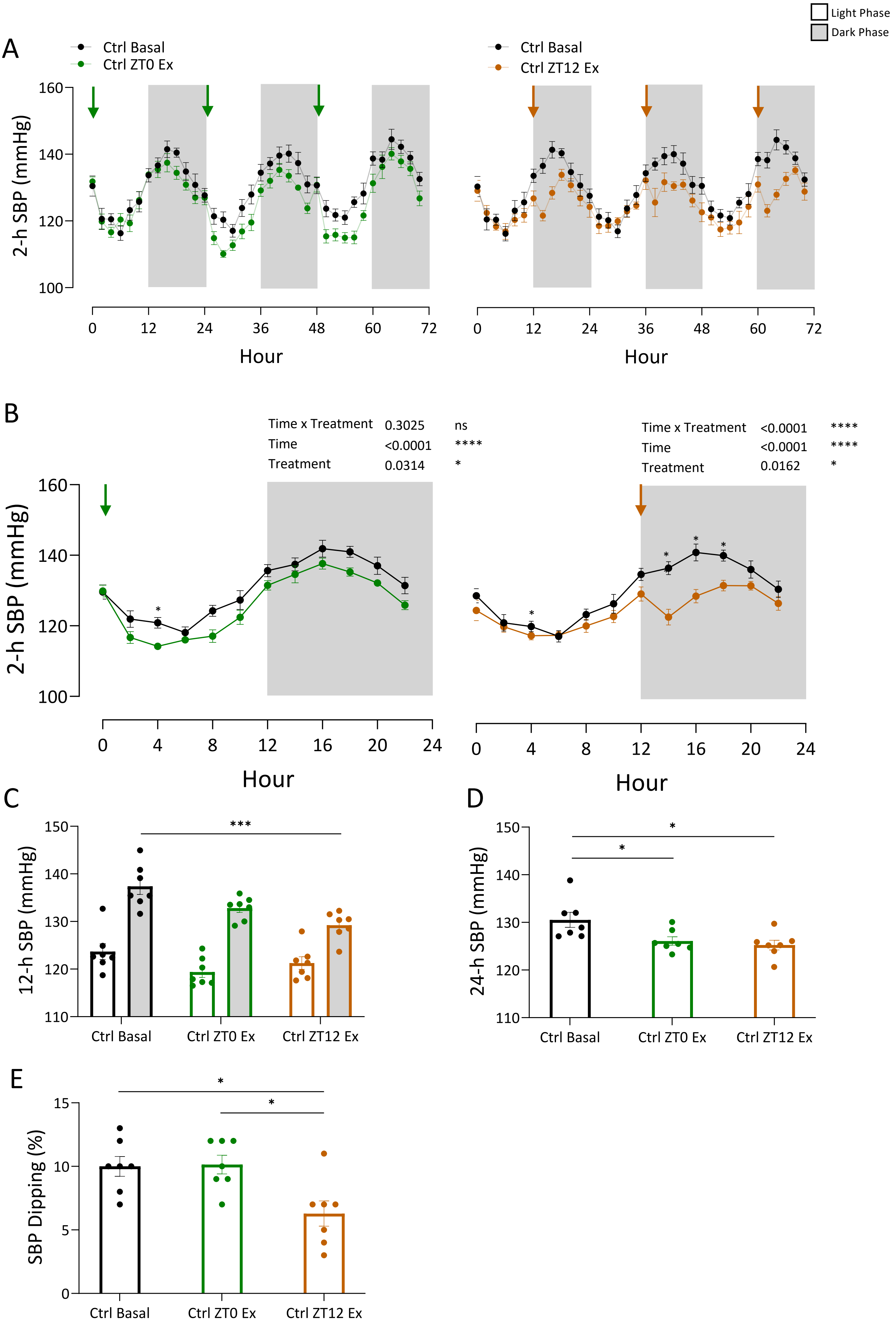


**Supplemental Figure S6. Exenatide administration results in a significant decrease in SBP in nondiabetic controls. [A]** 2-hour SBP bins collected over 24 hours under basal (black), after ZT0 exenatide administration (green), or ZT12 exenatide administration (orange). Arrows in green or orange indicate ZT0 or ZT12 injection of exenatide (20 μg/kg), respectively. **[B]** 2-hour SBP bins over 72-hours averaged to one 24-hour period. **[C]** 72-hours of SBP averaged to light/dark phases (12-h). **[D]** 72 hours of SBP averaged to 24 hours. **[E]** SBP dipping - the percentage of SBP decrease from dark to light phase. Data were analyzed by two-way ANOVA (B,C) and one-way ANOVA (D,E) with multiple comparisons test and were expressed as the mean ± SEM. *P < 0.05; **P < 0.01; ***P < 0.001, ****P < 0.0001; ns, not significant. Data were analyzed by two-way ANOVA with Sidak's (B) or Tukey's (C) multiple comparisons post-hoc analysis, and one-way ANOVA (D,E) with Tukey's multiple comparisons post-hoc analysis and were expressed as the mean ± SEM. *P < 0.05; **P < 0.01; ***P < 0.001, ****P < 0.0001; Data not indicated, or labeled as ns, is not significant.

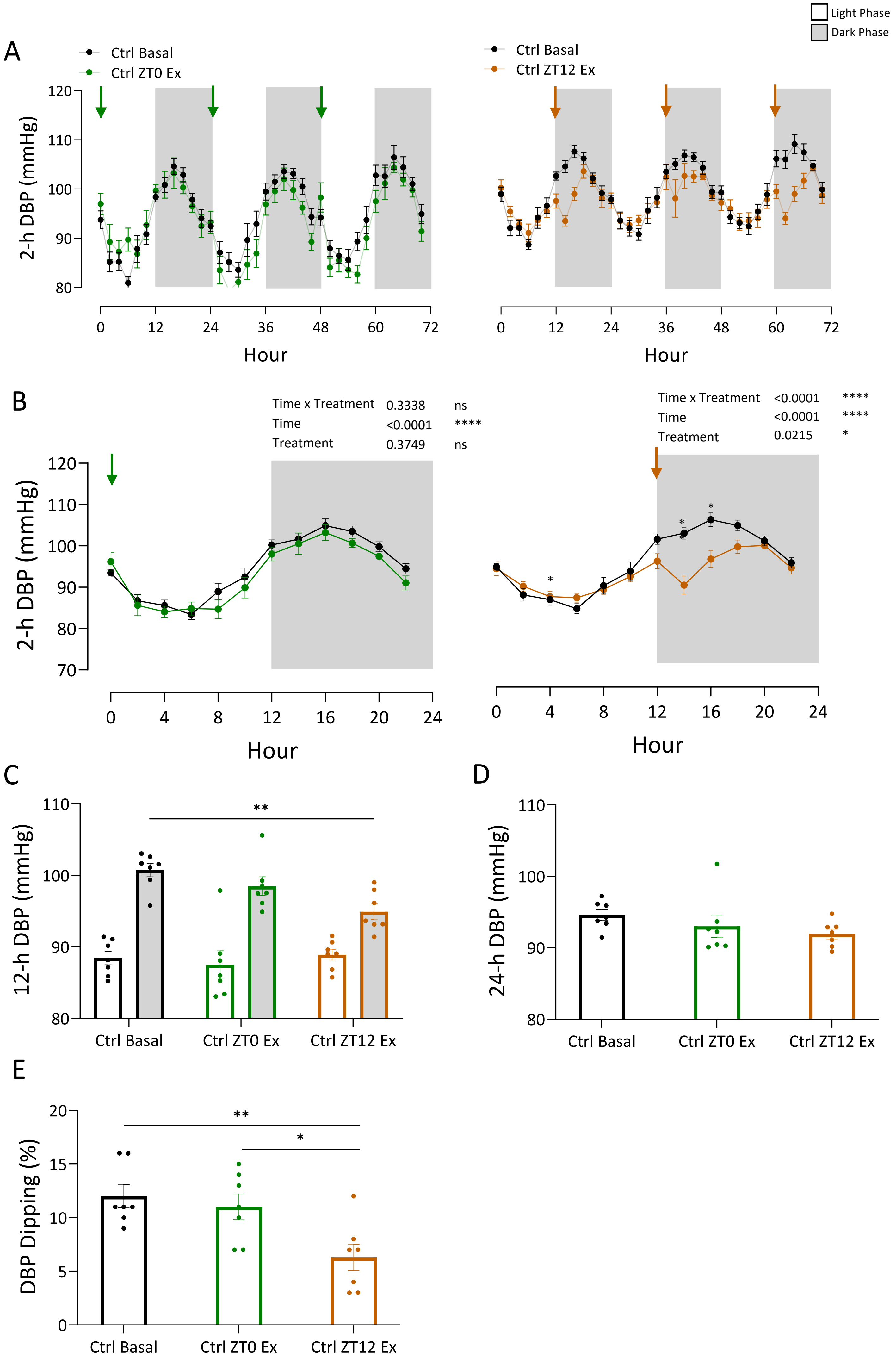


**Supplemental Figure S7. ZT12, but not ZT0, exenatide significantly decreases DBP in nondiabetic controls. [A]** 2-hour DBP bins collected over 72 hours under basal (black), after ZT0 exenatide administration (green), or ZT12 exenatide administration (orange). Arrows in green or orange indicate ZT0 or ZT12 injection of exenatide (20 μg/kg), respectively. **[B]** 2-hour DBP bins over 72-hours averaged to one 24-hour period. **[C]** 72 hours of DBP averaged to light/dark phases (12-h). **[D]** 72 hours of DBP averaged to 24 hours. **[E]** DBP dipping - the percentage of DBP decrease from dark to light phase. Data were analyzed by two-way ANOVA with Sidak's (B) or Tukey's (C) multiple comparisons post-hoc analysis, and one-way ANOVA (D,E) with Tukey's multiple comparisons post-hoc analysis and were expressed as the mean ± SEM. *P < 0.05; **P < 0.01; ***P < 0.001, ****P < 0.0001; Data not indicated, or labeled as ns, is not significant.

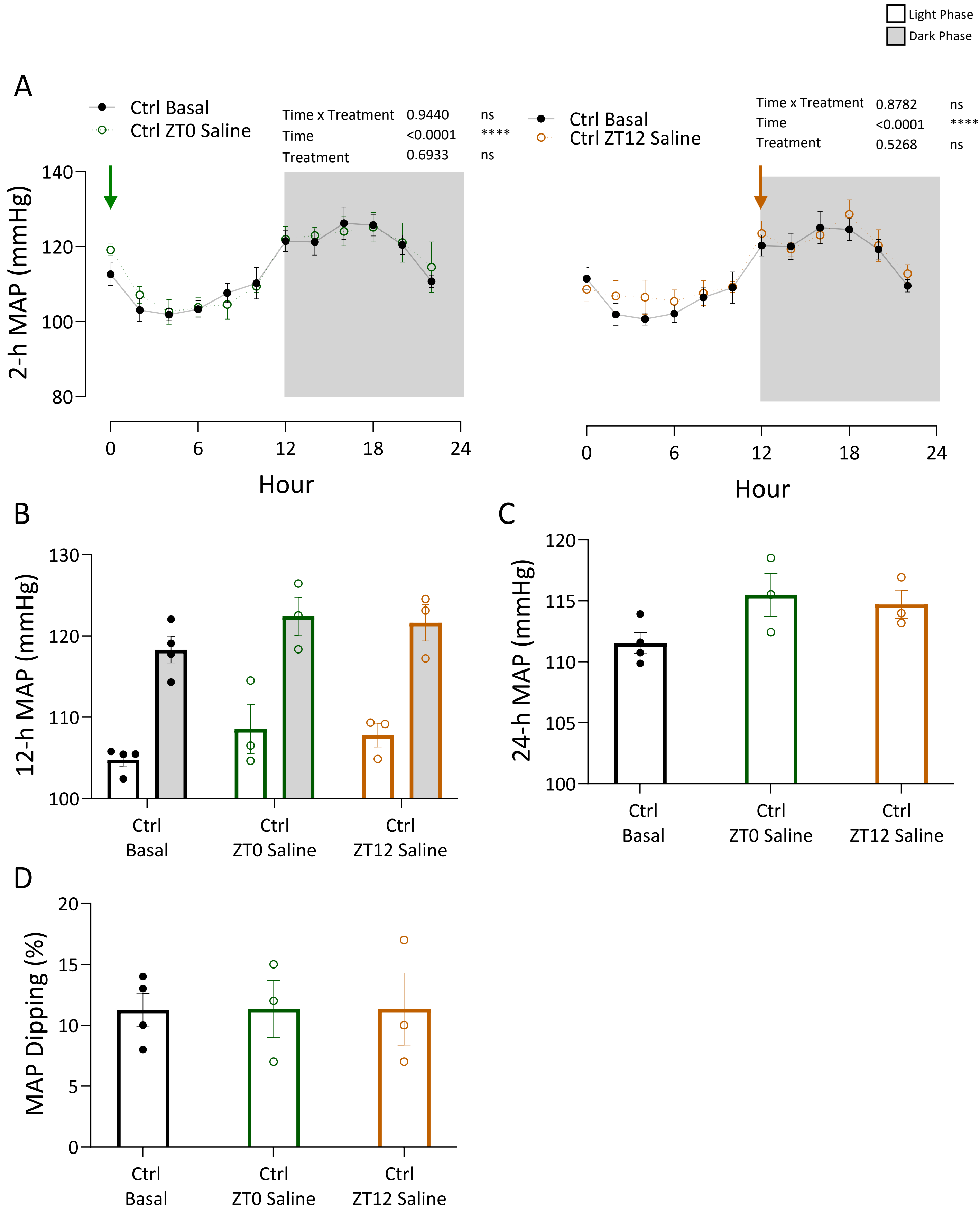


**Supplemental Figure S8**. **Neither ZT0 nor ZT12 saline administration reduces BP in nondiabetic control mice. [A]** 2-hour MAP bins collected over 24 hours under basal (black), after ZT0 exenatide administration (green), or ZT12 exenatide administration (orange). Arrows in green or orange indicate ZT0 or ZT12 injection of exenatide (20 µg/kg), respectively. **[B]** 12-hour bins of MAP **[C]** 24-hour average of MAP **[D]** MAP dipping - the percentage of MAP decrease from dark to light phase. Data were analyzed by two-way ANOVA with Sidak's (A) or Tukey's (B) multiple comparisons post-hoc analysis, and one-way ANOVA (C, D) with Tukey's multiple comparisons post-hoc analysis and were expressed as the mean ± SEM. *P < 0.05; **P < 0.01; ***P < 0.001, ****P < 0.0001; Data not indicated, or labeled as ns, is not significant.

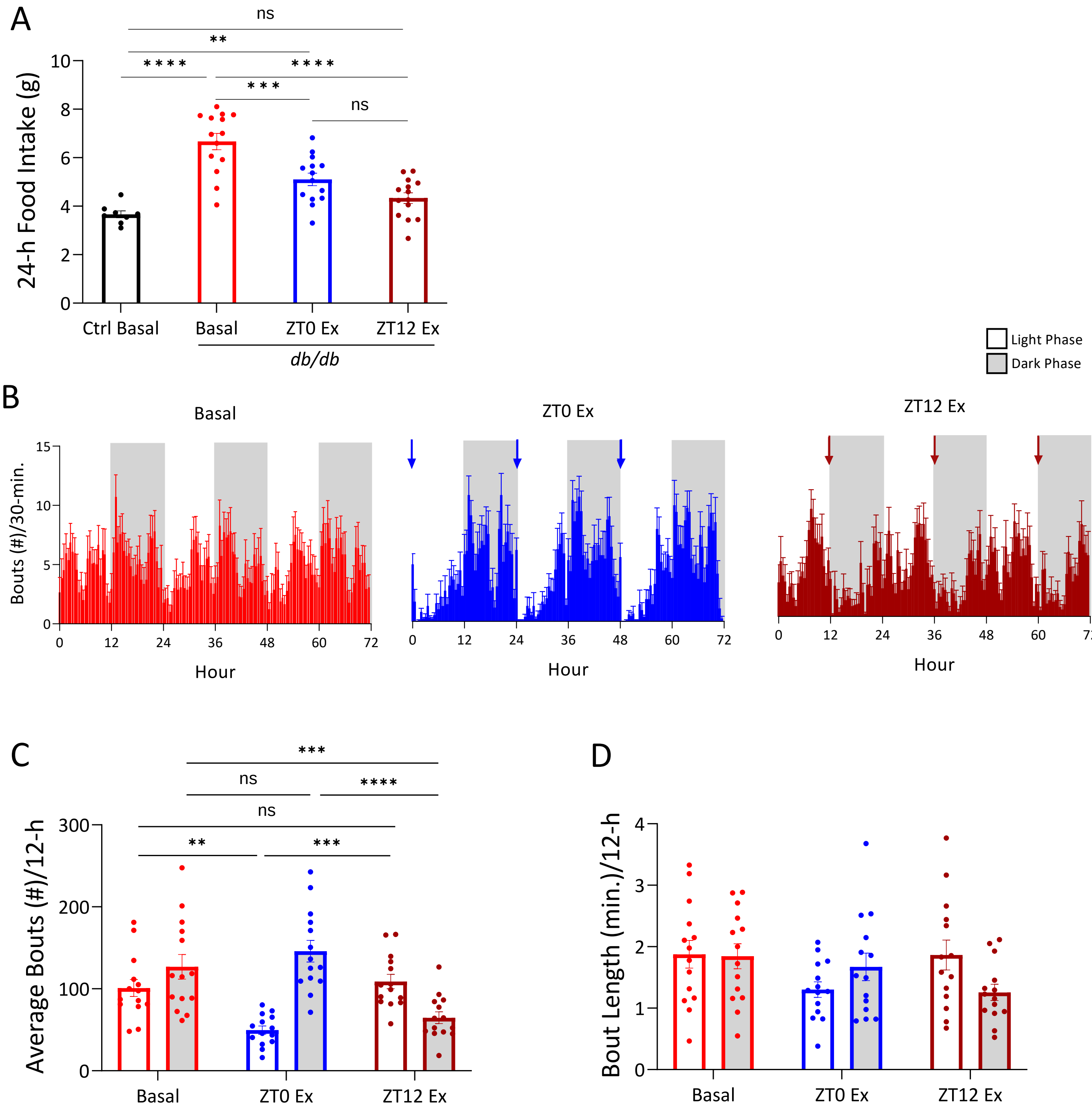


**Supplemental Figure S9. Food intake characteristics before and after exenatide treatment in *db/db* mice.** **[A]** 24-h average food intake. **[B]** The number of bouts in 30-min. intervals over 72-hours under basal (red), with ZT0 exenatide administration (blue), with ZT12 exenatide administration (dark red). Arrows in blue or dark red indicate ZT0 or ZT12 injection of exenatide (20 µg/kg), respectively. **[C]** 72-hours of bout number per 30-min. averaged to light/dark phases (12-h). **[D]** 72-hours of bout number per 30-min. averaged to 24-hours. Data were analyzed by two-way ANOVA (A,C,D) with Tukey's multiple comparisons post-hoc analysis, and were expressed as the mean ± SEM. *P < 0.05; **P < 0.01; ***P < 0.001, ****P < 0.0001; Data not indicated, or labeled as ns, is not significant.

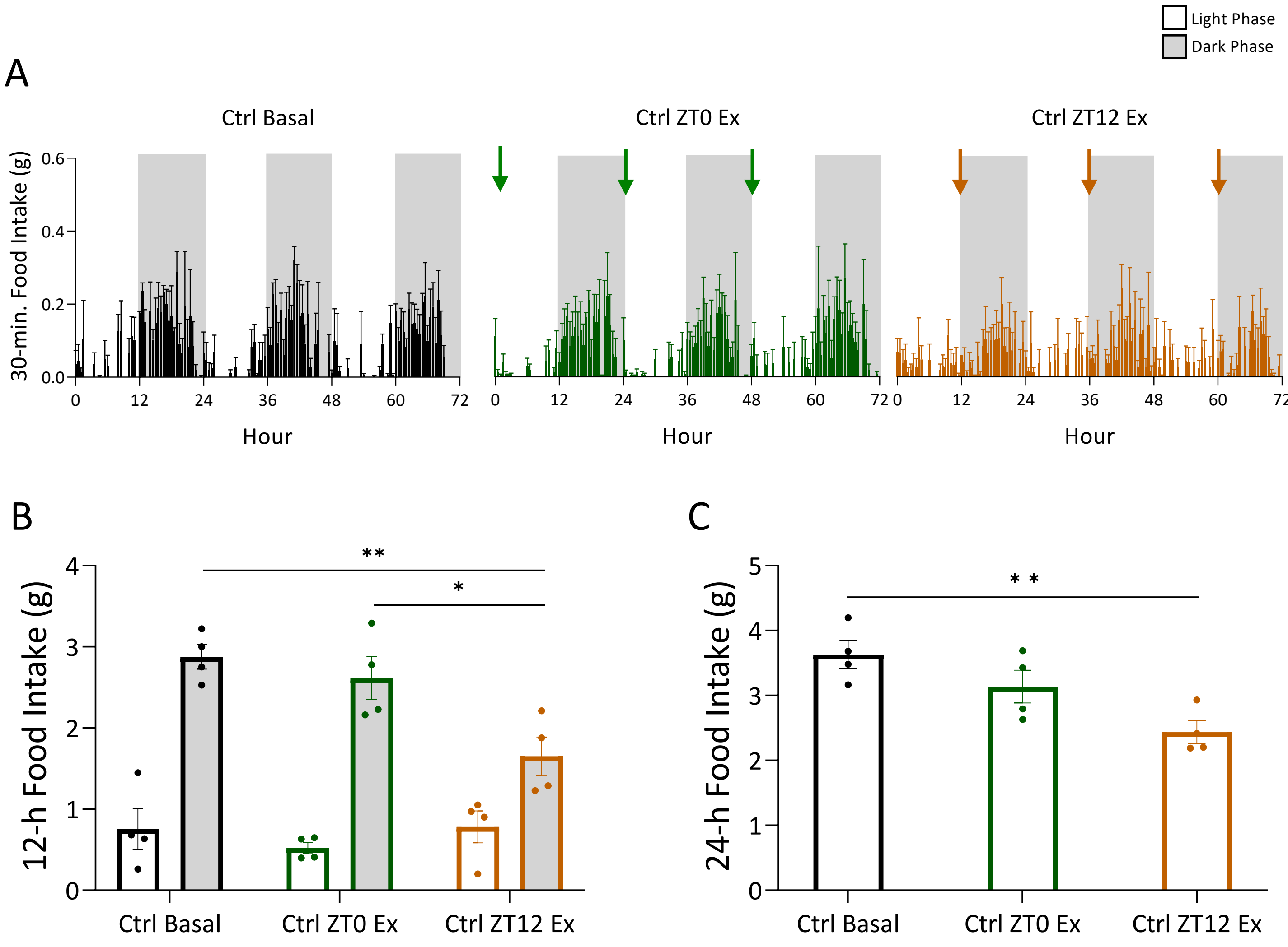


**Supplemental Figure S10. ZT12, but not ZT0 administration of exenatide suppressed food intake in nondiabetic control mice. [A]** 30-minute food intake bins over 72 hours under basal conditions (black), with ZT0 exenatide administration (green), and with ZT12 exenatide administration (orange). Arrows in green or orange indicate ZT0 or ZT12 injection of exenatide (20 µg/kg), respectively. **[B]** 72-hours of food intake averaged to light/dark phases (12-h). **[C]** 72-hours of food intake averaged to 24-hours. Data were analyzed by two-way ANOVA (B) or one-way ANOVA (C) with Tukey's multiple comparisons post-hoc analysis, and were expressed as the mean ± SEM. *P < 0.05; **P < 0.01; ***P < 0.001, ****P < 0.0001; Data not indicated, or labeled as ns, is not significant.

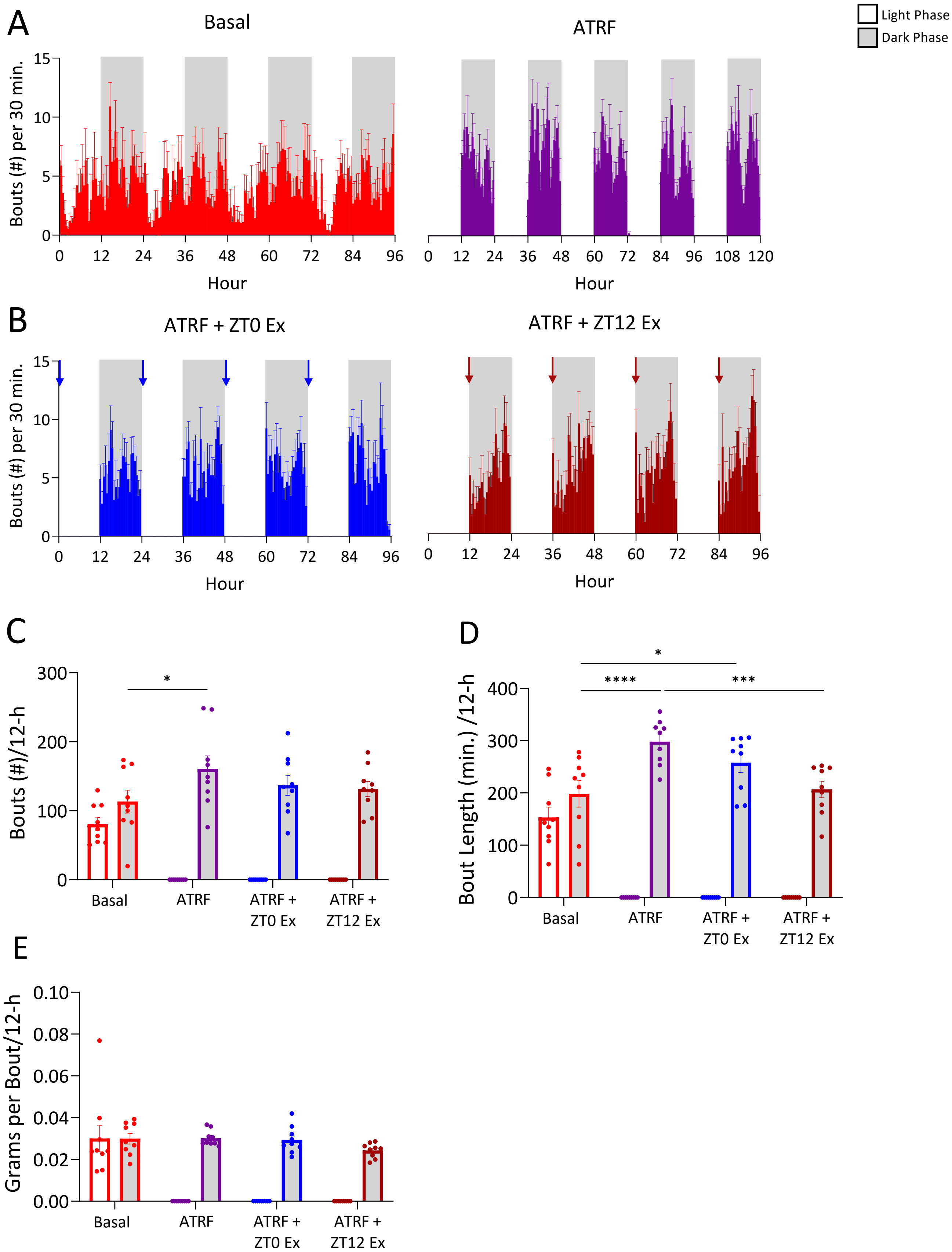


**Supplemental Figure S11. Reduction in food intake in *db/db* mice under ATRF and after exenatide administration is primarily the result of reduced bout length. [A]** The number of bouts in 30-min. intervals over time under basal (red) and under ATRF (purple) conditions. **[B]** The number of bouts in 30-min. intervals over time after ZT0 (blue) and ZT12 (dark red) exenatide administration. Arrows indicate time of injection. **[C]** Average number of bouts per light and dark phases (12-h). [D] Average length of bouts, in minutes, per light and dark phases. **[E]** Average amount of grams of food consumed per bout, per light and dark phases. Data were analyzed by two-way ANOVA (B,D,F) and one-way ANOVA (C,E,G) and with Tukey's multiple comparisons post-hoc analysis and were expressed as the mean ± SEM. *P < 0.05; **P < 0.01; ***P < 0.001, ****P < 0.0001; Data not indicated, or labeled as ns, is not significant.

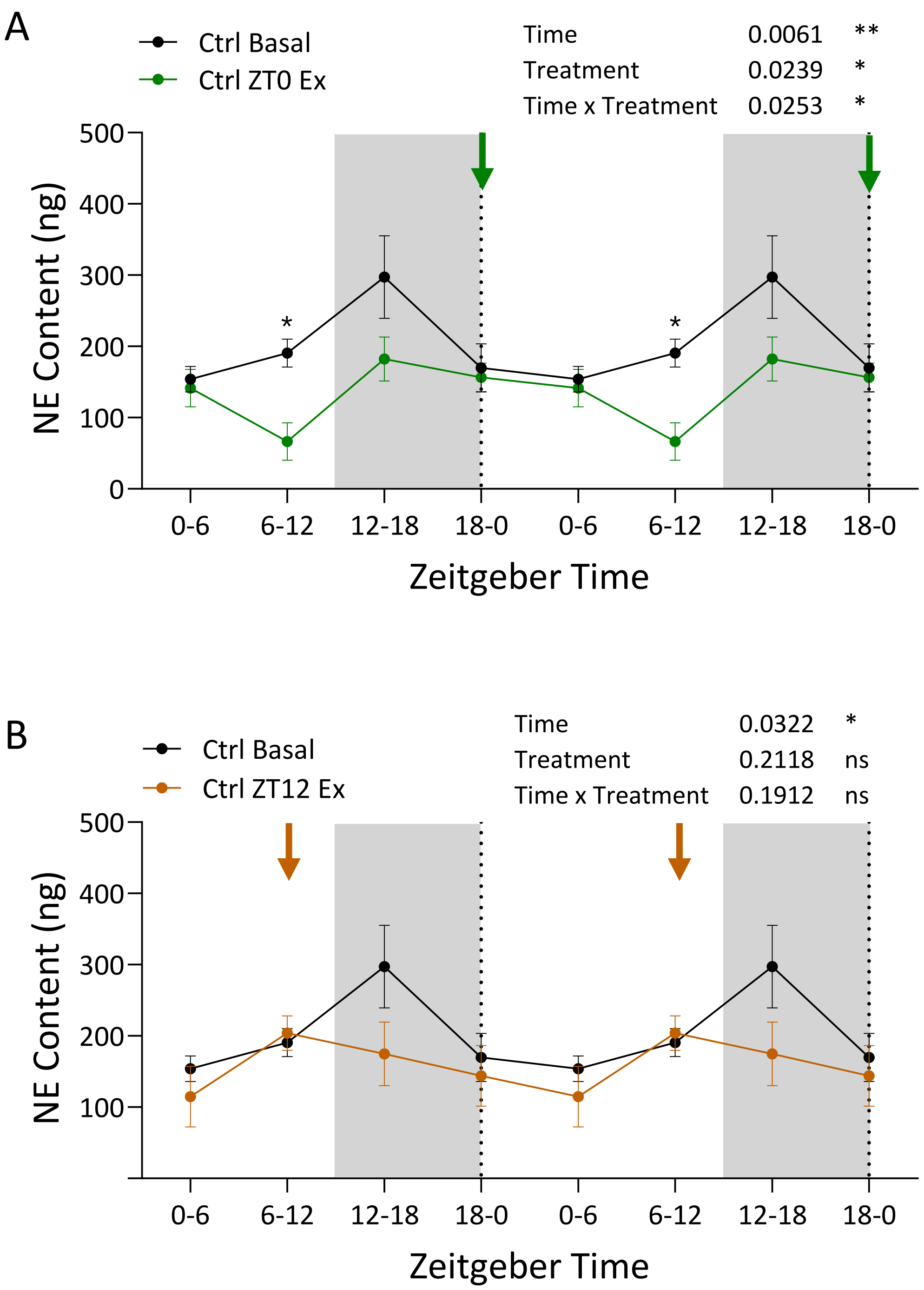


**Supplemental Figure 12. The effects of exenatide administration on NE content in nondiabetic control mice over 24 hours.** NE content in 6-hour urine samples was collected across a 24-hour day using HPLC. Urinary NE content of [A] control mice under basal conditions (black) and after ZT0 exenatide administration (green); and [B] control mice under basal conditions (black) and after ZT12 exenatide administration (orange). Levels of NE are expressed as total content, calculated by concentration (ng/mL) x urine volume (mL). Graphs were double-plotted to better illustrate the daily changes in NE content over time. Arrows indicate the time of exenatide injection. Data were analyzed by two-way ANOVA with Sidak's multiple comparisons post-hoc analysis and were expressed as the mean ± SEM. *P < 0.05; **P < 0.01; ***P < 0.001; Data not indicated, or labeled as ns, is not significant.
